# CYpHER: Catalytic extracellular targeted protein degradation with high potency and durable effect

**DOI:** 10.1101/2024.02.21.581471

**Authors:** Zachary R. Crook, Gregory P. Sevilla, Pamela Young, Emily J. Girard, Tinh-Doan Phi, Monique Howard, Jason Price, James M. Olson, Natalie W. Nairn

**Affiliations:** Cyclera Therapeutics Inc, Seattle, WA 98115, USA; Blaze Bioscience Inc., Seattle, WA 98109, USA; Clinical Research Division, Fred Hutchinson Cancer Center, Seattle, WA 98109, USA; Ben Towne Center for Childhood Cancer Research, Seattle Children’s Research Institute, Seattle, WA 98105, USA; NW Biosensor, Seattle, WA 98102

**Author notes:** Janssen Research and Development, Lower Gwynedd Township, PA 19002. Western University of Health Sciences, Lebanon, OR 97355. Present address of Z.R.C., G.P.S., and N.W.N.

## Abstract

Many disease-causing proteins have multiple pathogenic mechanisms, and conventional inhibitors struggle to reliably disrupt more than one. Targeted protein degradation (TPD) can eliminate the protein, and thus all its functions, by directing a cell’s protein turnover machinery towards it. Two established strategies either engage catalytic E3 ligases or drive uptake towards the endolysosomal pathway. Here we describe CYpHER (<u>C</u>atal<u>Y</u>tic <u>pH</u>-dependent <u>E</u>ndolysosomal delivery with <u>R</u>ecycling) technology with potency and durability from a novel catalytic mechanism that shares the specificity and straightforward modular design of endolysosomal uptake. By bestowing pH-dependent release on the target engager and using the rapid-cycling transferrin receptor as the uptake receptor, CYpHER induces endolysosomal target delivery while re-using drug, potentially yielding increased potency and reduced off-target tissue exposure risks. The TfR-based approach allows targeting to tumors that overexpress this receptor and offers the potential for transport to the CNS. CYpHER function was demonstrated *in vitro* with EGFR and PD-L1, and *in vivo* with EGFR in a model of EGFR-driven non-small cell lung cancer.

## Introduction

Contemporary targeted therapeutics aim to modulate the activity of a particular target, usually a protein, that has a defined role in disease pathology. This modulation is often the disruption of protein function, most commonly seen by enzyme inhibition (e.g., kinase inhibitors) or steric blocking (e.g., antibodies). These conventional inhibitors and blockers can disrupt a defined function, and often come with beneficial pleiotropic effects, such as induction of protein homeostasis disruption *(1)* or altered target trafficking *(2, 3)*. Nevertheless, these drugs are often insufficient to meaningfully and durably alter disease pathology. For one, many targets exhibit multiple functions, and inhibiting one function can leave the others available for potentiating pathologic signaling. Second, the nature of many of these inhibitors leaves them particularly vulnerable to cellular adaptation and mutational resistance that diminishes drug durability.

An exemplary group of protein targets is receptor tyrosine kinases (RTKs) in cancer, as they can potentiate growth, differentiation, and survival signaling *(4)*. Many involve a mechanism for activation that involves multimerization and cross-phosphorylation at the cell surface. As such, they not only function as kinases, but as kinase substrates mediating signal transduction. Tyrosine kinase inhibitors (TKIs) and antibodies are typical RTK targeted therapeutics. TKIs can disrupt kinase activity of RTKs *(5)*, but they do not block the RTKs from being substrates for other kinases. As many RTKs function through both homo- and heterodimerization with other RTKs, this leads to resistance via upregulation of other partners *(6, 7)*. As an additional liability, point mutations can often directly or indirectly disrupt TKI binding *(7, 8)*. Conversely, antibodies can alter the multimerization tendencies of RTKs, but with kinase function retained, target or heterodimer partner upregulation (or gain-of-function mutation) becomes a common resistance mechanism *(9)*, with increased total membrane kinase activity compensating for disrupted multimerization. Altogether, these inhibitors can be effective for a period of time, but the myriad mechanisms for functional bypass typically render their efficacy transient.

In order to simultaneously disrupt all of a target’s disease-associated functions, targeted protein degradation (TPD) can be used. Expertly summarized in a recent review *(10)*, TPD leverages cells’ mechanisms for target turnover, altering the kinetics of this process for specific targets. This is mainly through formation of ternary complexes between target protein and degradation effector. For intracellular (e.g., molecular glue and PROTAC) *(11)* and some extracellular (e.g., AbTAC and REULR) *(12, 13)* molecules, this is through E3 ligase recruitment, inducing ubiquitin-mediated target degradation by the proteasome or lysosome. This benefits from a catalytic mechanism of action (the drug is not expended in the process) but can be complicated by finding an E3 ligase expressed in the target tissue that can be induced to interact in an effective orientation with the target for functional ubiquitination.

Meanwhile, small molecules (e.g., molecular glue and PROTAC) also have many of the same point mutational resistance liabilities of TKIs. Other approaches used for surface and extracellular soluble targets (e.g., LYTAC, ATAC, KineTAC) *(14–16)* engage surface protein trafficking systems. Often targeting membrane sugar receptors (cation-independent mannose-6-phosphate receptor [CI-M6PR] or asialoglycoprotein receptor [ASGPR]) but also cytokine receptors (e.g., CXCR7), these are designed to “hitch a ride” with the uptake receptor into the cell before target release in the endolysosomal system. These tend to be biologics, benefiting from modular design and greater target specificity, but the drug follows the target through its trafficking, limited to stoichiometric (as opposed to catalytic) activity.

We have developed an extracellular TPD (eTPD) technology, combining the specificity and modularity of an endolysosomal trafficking approach with a unique catalytic mechanism, that we call CYpHER (**C**atal**Y**tic **pH**-dependent **E**ndolysosomal delivery with **R**ecycling). Here, we engage a rapidly-recycling uptake receptor, transferrin receptor (TfR). Its normal role is facilitating uptake of iron-loaded transferrin *(17)*. Upon uptake and endosomal maturation, which involves acidification to roughly pH 5.5, transferrin releases its iron but remains bound to TfR, returning to the surface with its receptor. The whole process takes ∼10-20 mins *(17, 18)*, repeating dozens to hundreds of times over the protein’s lifetime (with a natural turnover half-life of ∼24 hrs) *(18, 19)*. CYpHER molecules mimic the behavior of transferrin, except the cargo is a target for elimination. The target-binding end of CYpHER, using existing or engineered properties, has reduced binding affinity at endosomal pH, permitting target release and subsequent trafficking through the endolysosomal system. Conversely, its TfR-binding end has no such pH sensitivity, permitting the CYpHER to return to the surface to take in additional target molecules. Catalytic activity can increase potency (multiple target molecules eliminated by a single drug), permits retained activity after drug is cleared from extracellular space to increase the durability of function, and reduces the disruptive effects of shed, soluble variants of a membrane target (that can otherwise act as a decoy for conventional antibodies) *(20, 21)*, since the soluble form will simply represent one round of uptake and endosomal release.

Using TfR as the uptake receptor presents numerous advantages, particularly for CNS disease and oncology. TfR is commonly upregulated on a wide variety of solid tumors *(22, 23)*, presumably to accommodate the increased iron demands of energy generation and nucleotide synthesis in rapidly-dividing cells *(24)*, and its overexpression often correlates with disease severity *(23, 25, 26)*. This overexpression compared to healthy tissue should concentrate the drug in the tumor tissue, sparing other tissue from potential toxicities and increasing the therapeutic window. Also, unlike other uptake receptors, cancer cells are highly dependent on TfR for growth and survival (based on genome-wide knockout data from >1000 cancer cell lines) *(27)*, so a potential resistance mechanism of reduced receptor expression is avoided by using TfR. In addition, TfR is well known as a mediator of blood-brain barrier transcytosis *(28)*, having been used to deliver biologic molecules to the brain parenchyma *(29, 30)*. A drug whose mechanism of action includes TfR engagement has the potential to enable specific depletion of targets in the CNS, an area of high unmet medical need. We will note that, during the preparation of this work, another group also highlighted the advantages of TfR-based TPD for oncology (TransTACs) *(31)*, but designed the molecules to be degraded in the endosome to avoid (rather than harness) TfR-based recycling for catalytic activity.

Here we present our work on developing molecules with these properties, demonstrating catalytic target uptake and elimination, suppressing signaling and growth of cancer cells, and *in vivo* pharmacokinetics and activity. We discuss the steps for engineering such molecules with these characteristics and assays for demonstrating surface target elimination and uptake of multiple targets per drug molecule, and then discuss potential advantages and utilities of this approach.

## Results

### The CYpHER concept

The core CYpHER concept is illustrated in Fig. 1, A and B. A simplistic diagram of a CYpHER molecule (Fig. 1A) includes a pH-dependent target-binder linked to a recycling uptake receptor-binder (here, a TfR binder). After CYpHER has initiated ternary complex formation on the cell surface (Fig. 1B), the target and CYpHER molecule are brought into the cell in TfR-containing clathrin-coated vesicles. Following the standard trafficking of TfR, the vesicle then joins endosomes that begin acidifying *(32)*. The target-binders are engineered to release the target at low pH, permitting the TfR:CYpHER complex to be trafficked back to the surface where the target-binding end of CYpHER is free to bind and traffic additional target molecules. Meanwhile, the released target undergoes intracellular trafficking which include lysosomal delivery and subsequent degradation.

**Fig. 1.**
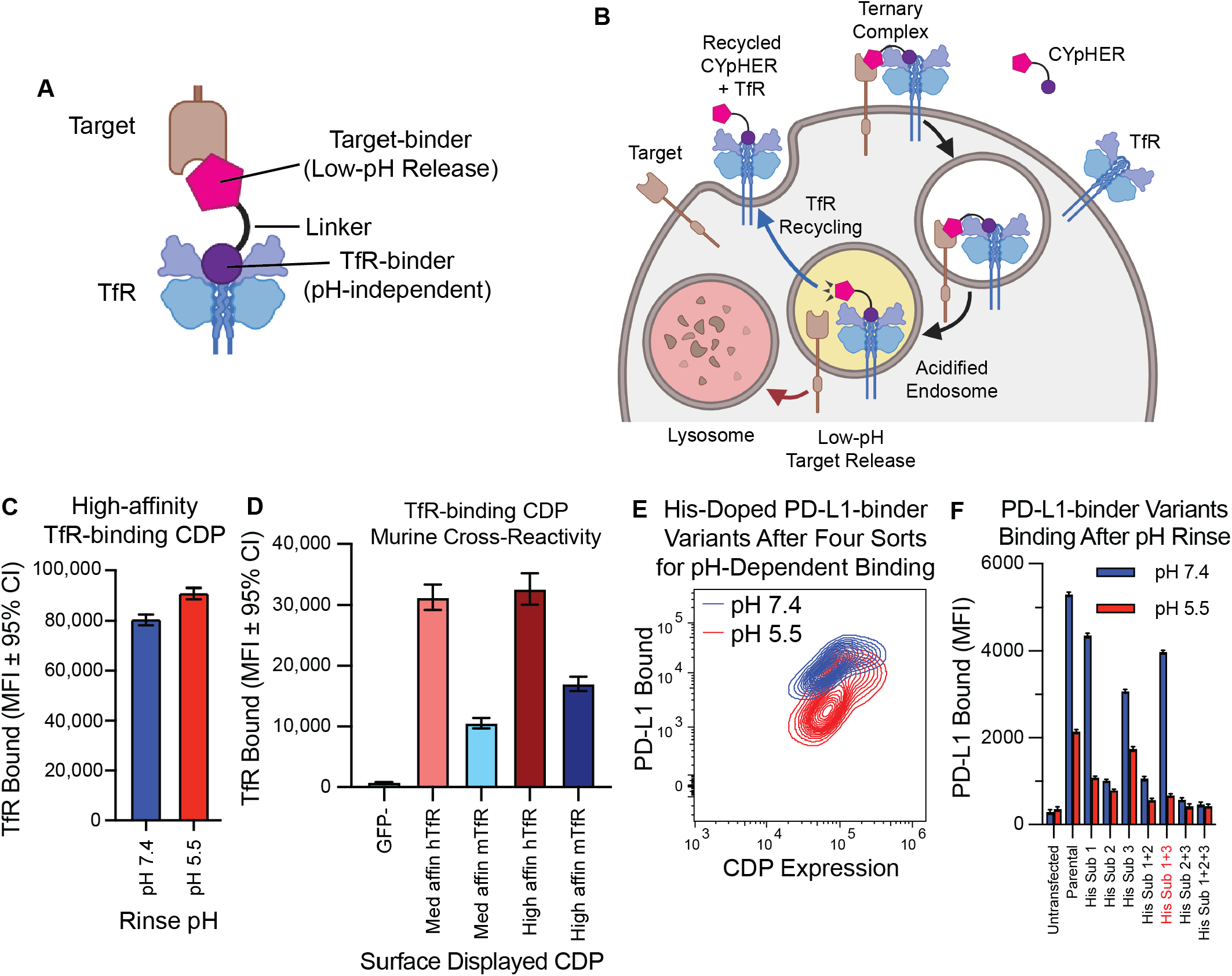
Basic principles of CYpHER and component binders. (**A**) CYpHER design including a pH-independent TfR-binding domain and a pH-dependent target-binding domain separated by a linker. (**B**) CYpHER mechanism. CYpHER induces ternary complex formation with target and TfR. Upon TfR-mediated uptake and endosomal acidification, target is released for endolysosomal system trafficking. TfR and CYpHER recycle to the surface for engagement with another target molecule. (**C**) 293F cells displaying a high-affinity TfR-binding CDP were stained with TfR and rinsed at pH 7.4 or pH 5.5 for 10 minutes, showing similar binding in both conditions. (**D**) 293F cells displaying medium or high affinity TfR-binding CDPs were stained with human TfR (hTfR) or mouse TfR (mTfR). (**E**) pH-dependent PD-L1 binding flow profile of 293F cells displaying a pool of histidine-doped variants of a PD-L1-binding CDP after four rounds of flow sorting; two for high binding after pH 7.4 rinse, two for low binding after pH 5.5 rinse. (**F**) Three His substitutions were tested as singletons and combinations for PD-L1-binding after 10 minutes pH 7.4 or pH 5.5 rinse. Variant with His substitutions 1 and 3 was chosen for further work.

To construct CYpHER molecules, we began with two cystine-dense peptide (CDP) miniproteins that we’ve recently characterized separately for other *in vivo* functionalities *(30, 33)*. We made use of a TfR-binding CDP allelic series *(30)* that maintains binding at reduced pH (Fig. 1C), has murine cross-reactivity (Fig. 1D) as demonstrated in mammalian surface display staining experiments, and also has access to the CNS *(30)*. Then we further engineered a PD-L1-binding CDP *(33)* for enhanced pH-dependent release by generating a library of variants that each contained up to two histidine substitutions, and surface-display screened to enrich for variants that conferred strong PD-L1 binding at pH 7.4 but reduced PD-L1 binding to cells after a low-pH rinse (Fig. 1E). Three singleton histidine substitutions were significantly enriched, and when testing them as singletons and combinations, one combination (His Sub 1+3) was found to confer PD-L1-binding almost as well as the parental variant at pH 7.4 but conferred substantially less PD-L1 binding after a low-pH rinse (Fig. 1F).

To confirm cooperative target and TfR binding, a pilot molecule was produced that consisted of the pH-dependent PD-L1-binding CDP fused to a high affinity member of the TfR-binding CDP series via a flexible Gly-Ser linker (fig. S1A). 293T cells were transfected with PD-L1-GFP and TfR-RFP (or with TfR-RFP alone) and then incubated with 10 nM pilot molecule for 24 hr. Subsequent cell staining for a 6xHis tag on the pilot molecule and gating for double negative, GFP+/RFP-, GFP-/RFP+, or GFP+/RFP+ cells indicated increased cell staining when one or the other surface binding partner is overexpressed with a massive increase in cell staining when both are overexpressed (24-42x vs one or the other) (fig. S1B). This confirmed the cooperative targeting that could be achieved.

### Engineering CYpHER candidates

Further CYpHER candidates were generated that included Fc domains for increased *in vivo* serum exposure (Fig. 2A). Both candidates contained an Fc domain with a high-affinity TfR binder fused to its C-terminus by a flexible Gly-Ser linker. One candidate (CT-4212-1) also contained an N-terminal fusion (via flexible Gly-Ser linker) with the pH-dependent PD-L1-binding CDP, while the other candidate (CT-4212-3) used a rigid linker derived from human IgA to fuse the PD-L1-binding CDP to the C-terminus of the TfR binder. Both molecules were tested on polyclonal 293T, H1650, and MDA-MB-231 cell populations transduced (via lentivirus) to express PD-L1-GFP. A functional CYpHER is expected to drive PD-L1-GFP from the cell surface (Fig. 2B). In all three populations (Fig. 2, C to E), PD-L1 was trafficked from the surface, as seen in both microscopy and flow cytometry assays (the latter measured by staining with a non-competitive PD-L1 antibody [clone 22C3]) *(34)*. Flow cytometric quantitation of per-cell GFP level changes demonstrated substantial and consistent surface PD-L1 removal (Fig. 2, F, H, and J), as well as overall PD-L1-GFP reduction in all cases (Fig. 2, G, I, and K); the degree of total PD-L1-GFP reduction varied by cell line and CYpHER. The highest-expressing polyclonal population, 293T-PDL1-GFP, experienced the largest degree of GFP reduction. This could be caused by better cooperative CYpHER accumulation when more target is present, by saturation of the recycling pathway, or both.

**Fig. 2.**
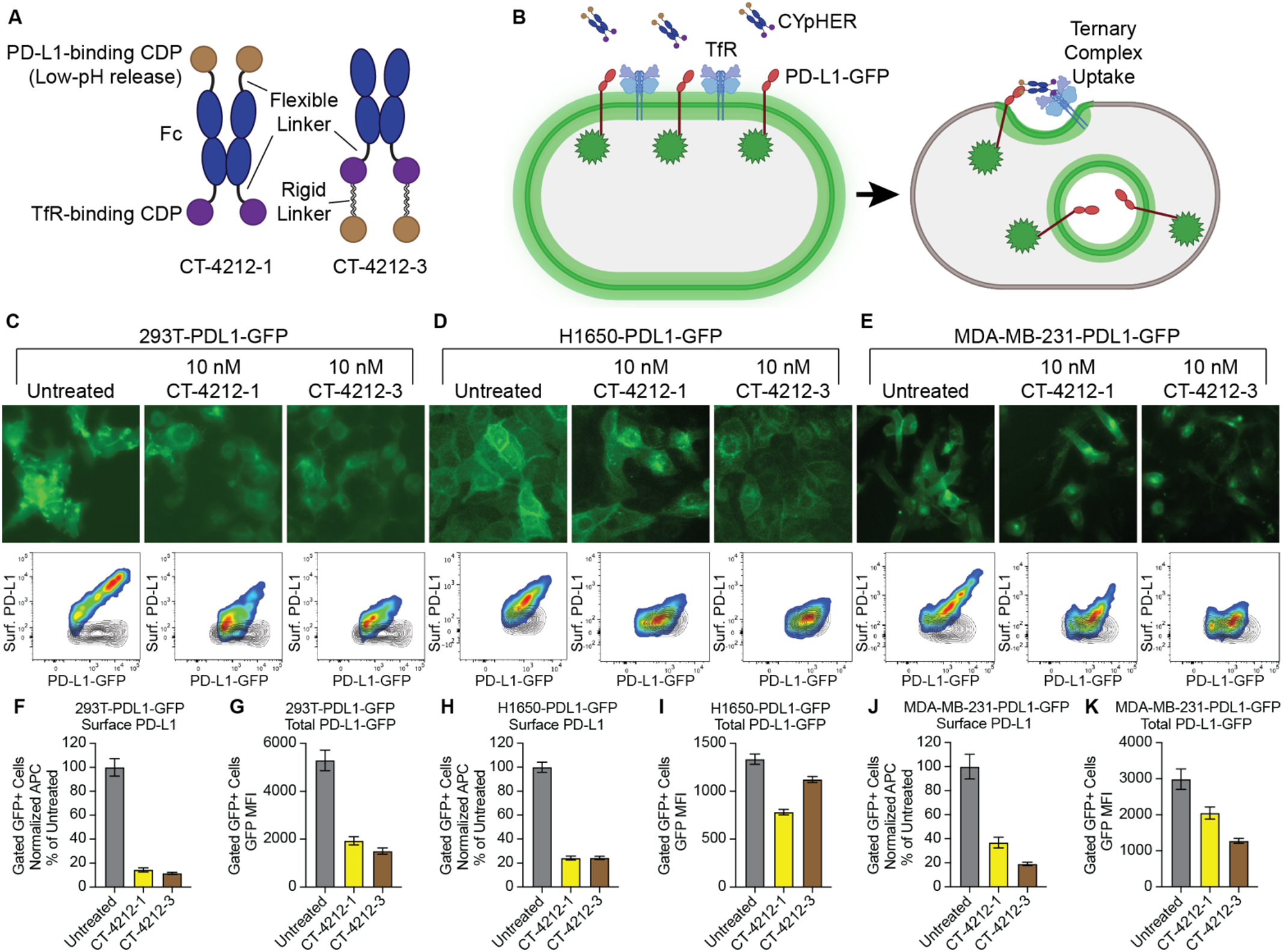
PD-L1 CYpHER design and target depletion in cell pools overexpressing PD-L1-GFP. (**A**) Two designs of PD-L1 CYpHERs, named CT-4212-1 and CT-4212-3, using a high-affinity TfR-binding CDP and a pH-dependent PD-L1-binding CDP. (**B**) Illustration of PD-L1-GFP trafficking induced by CYpHER. (**C** to **E**) Pools of 293T (**C**), H1650 (**D**), and MDA-MB-231 (**E**) cells transduced with lentivirus driving PD-L1-GFP were untreated or incubated with 10 nM CYpHER for 24 hr before GFP-channel microscopy (above) and flow cytometry (below) after staining for surface PD-L1. Black contour in flow profiles: cells stained without PD-L1 antibody. (**F** to **K**) Quantitation of normalized surface PD-L1 (**F, H**, and **J**) or total PD-L1-GFP (**G, I**, and **K**) signal in 293T-PDL1-GFP (**F** and **G**), H1650-PDL1-GFP (**H** and **I**), and MDA-MB-231-PDL1-GFP (**J** and **K**) cells with or without CYpHER treatment.

The CYpHER platform is amenable to any target for which surface or soluble elimination would provide benefits to patients beyond simple binding or blocking of one site. One such target, with dual roles as both kinase and kinase substrate (as well as other cell-surface-driven functions) *(35)*, is EGFR. It is implicated as a primary driver of several of the most common and deadly cancers, including glioblastoma, lung cancer, head and neck cancer, and colon cancer *(6, 36–39)*. It is primarily targeted by TKIs that disrupt its kinase activity or monoclonal antibodies (mAbs) that block ligand-binding and subsequent homodimerization, the latter of which can also induce degradation by engaging natural ubiquitination pathways. Patients treated with these targeted molecules can benefit for a time, but resistance inevitably emerges and remains an area of urgent need for new therapies *(40)*. These resistance mechanisms are often either a point mutation to reduce TKI activity or an upregulation of another EGFR heterodimerization partner like HER2, ERBB3, or MET. Such upregulations can lead to cross-phosphorylation of EGFR and other RTKs, a process that is insufficiently suppressed by mAbs vs EGFR. Elimination of EGFR from the surface would drastically reduce such signaling, as the majority of oncogenic EGFR signaling, including kinase-independent signaling *(41–43)*, occurs at the plasma membrane.

An EGFR-binding VHH nanobody *(44)* was engineered into a CYpHER component through similar means to the PD-L1-binding CDP. In one method, a pool of variants with His substitutions was screened in mammalian surface display *(45, 46)* through four rounds of enrichment; two rounds enriched for high binding at neutral pH (pH 7.4), while the other two enriched for low binding at early endosomal pH (pH 5.5) (fig. S2A). For comparison, the final round of sorting also collected the population with high binding at pH 5.5 (fig. S2B). The primary variant in the pool enriched for low binding in low pH loses roughly half its bound EGFR in tetravalent (streptavidin) stain conditions upon low-pH rinse (fig. S2C), while the dominant variant from the population with high binding at low pH does not have this property (fig. S2D). Fc fusions to each of these molecules, when used as ligands in surface plasmon resonance experiments, verified these properties (fig. S2, E and F); the variant engineered for pH-dependent EGFR release demonstrated higher affinity at neutral pH (*K*_D_ = 16.2 ± 0.1 nM at pH 7.4) compared to low pH (*K*_D_ = 61.2 ± 0.3 nM at pH 5.8) and the other variant demonstrating slightly lower affinity at neutral pH (*K*_D_ = 10.5 ± 0.1 nM at pH 7.4) compared to low pH (*K*_D_ = 7.8 ± 0.07 nM at pH 5.8). Both of these represent >10-fold higher affinity vs the reported affinity of the parental nanobody *(47)*. The pH-dependent release nanobody was named EGFR Nanobody v1 and was incorporated into various CYpHER designs.

As an additional allele identification strategy, singleton His scanning (i.e. single His substitutions tested one at a time) was performed on the nanobody in CDRs 1 and 3, as these CDRs are primarily implicated in EGFR-binding *(47)*. One variant (His Sub 10) was identified in mammalian surface display that lost roughly half its bound EGFR in tetravalent stain conditions upon low-pH rinse (fig. S2G); this was named EGFR Nanobody v2.

### Adapting a native ligand for CYpHER

In building the platform, it was apparent that the target-binding domain needn’t be an exogenous molecule. As a great many disease-associated target proteins do so by signal transduction mediated by ligand-binding, the ligand presents itself as a natural target-binder that can be adapted, through engineering and affinity / pH maturation, for CYpHER incorporation. In the case of EGFR, EGF itself (a naturally pH-dependent binder which is also a CDP) *(48)* can be used after engineering it to disable signal transduction capabilities; here, binding to both Domain I and Domain III of EGFR induces a conformational change that renders the dimerization domain (Domain II) solvent-accessible *(49)*. This facilitates homo- or heterodimerization partner cross-phosphorylation in the cytosolic kinase domains and subsequent signal transduction through pathways that include MAPK, PI3K/AKT, PLC/PKC, and JAK/STAT *(50)*. Engineering EGF to eliminate Domain I-binding while enhancing Domain III-binding would produce a dominant-negative EGF variant, and candidate CYpHER component, that competitively engages EGFR but does not induce Domain II-dependent dimerization and signal transduction.

Producing such a variant began with identifying an EGF variant that binds to isolated Domain III, commercially available in the form of soluble EGFRvIII, a constitutively-active splice isoform of particular prominence in glioblastoma that is missing Domain I and part of Domain II *(51)*. EGF does not appreciably interact with EGFRvIII *(52)*, so we used Rosetta protein design software *(53–55)* to design and screen 488 variants that are predicted to have improved binding to Domain III based on a published co-crystal structure *(49)*. Mammalian surface display screening for binding to EGFRvIII produced two variants (fig. S3, A and B) named EGFd1 and EGFd2; EGFd1 demonstrated the strongest binding in surface display. The EGF:EGFR co-crystal structure *(49)* was further studied to identify four residues (M21, A30, I38, and W49) on EGF that contact Domain I such that mutations to them would be predicted to disrupt the interface, either eliminating hydrophobic interactions or introducing steric hindrance. Mutations to disrupt these residues one at a time (EGFd1.1 through 1.4), and one that disrupted all four at once (EGFd1.5), were tested for the ability to bind full-length EGFR or EGFRvIII. Three of the four Domain I interface point mutants (EGFd1.1, EGFd1.2, and EGFd1.3) and the quadruple mutant (EGFd1.5) demonstrated improved EGFRvIII binding, while all five EGFd1 variants demonstrated a substantial increase in the ratio of EGFRvIII binding vs full-length EGFR binding, indicating an apparent negative contribution of Domain I to EGFR binding (fig. S3C). Advancing EGFd1.5, it demonstrated a reduced stain in mammalian surface display upon pH 5.5 rinse, similar to that seen for EGFR Nanobody v1 (fig. S3D); as EGF is known to naturally possess pH-dependent EGFR binding *(48)*, this EGFd1.5 behavior confirmed the engineered variant retained this property. An additional round of affinity maturation (site-saturation mutagenesis followed by pooled mammalian display enrichment screening over two rounds of sorting) *(45)* yielded a higher-affinity variant (fig. S3E), EGFd1.5.36, that was chosen for testing in the CYpHER context.

### EGFR CYpHER induces EGFR surface clearance and elimination

A candidate EGFR CYpHER, CT-1212-1, was produced using a high-affinity TfR-binding CDP and EGFR Nanobody v1 (Fig. 3, A to C). The molecule demonstrated good expression and assembly with negligible aggregate after capture from supernatant and buffer exchange. 293T cells expressing a variable level of EGFR-GFP, expected to undergo target internalization upon CYpHER treatment similar to PD-L1-GFP (Fig. 3D), were dosed and analyzed by microscopy and flow cytometry (staining with non-competitive anti-EGFR clone 199.12) *(56)* for total EGFR-GFP and surface EGFR (Fig. 3E). As a whole population, cells demonstrated substantial (>80%) reduction of surface EGFR and ∼50% reduction of total EGFR-GFP signal, which was validated by Western blot of lysate from flow-sorted, viable cells (Fig. 3F). Flow sorting for viable cells was performed because dead cells and debris are not metabolically active and cannot drive endolysosomal trafficking of target proteins. Inspecting the flow profile, it was not “shifted” en-masse by CYpHER treatment; instead, much of the reduction occurred in those cells with the highest initial EGFR levels. Partitioning flow profiles from each treatment by surface EGFR supported this observation (Fig. 3G), where the cells with the most surface EGFR experienced the greatest proportional loss. This is further evidence for a possible mechanism involving either increased accumulation, saturable recycling pathways, or both; regarding the latter, this experiment used a clonal cell population that only differs by variable EGFR expression, so trafficking capabilities and limitations should be similar in all cells. Meanwhile, time-course experiments demonstrated that surface EGFR clearance is rapid (near maximal effect after 1 hr), consistent with rapid TfR-mediated uptake, while EGFR-GFP signal loss takes more time, likely due to the slower kinetics of lysosomal degradation vs surface internalization (Fig. 3, H and I). This was corroborated by labeling 293T-EGFR-GFP cells with LAMP1-RFP via baculovirus (BacMam 2.0, Invitrogen) for 24 hr and then treating cells with 10 nM DyLight 755-conjugated CT-1212-1 for 0, 1, or 4 hrs (Fig. 3J). Lysosomal RFP signal overlapped with EGFR-GFP foci more often and more intensely at 4 hr than at 1 hr. CYpHER signal was also more intense overall at 4 hr than at 1 hr. The CYpHER signal at 4 hr was within several intracellular compartments, including the lysosome, indicating some of the CYpHER being trafficked alongside EGFR-GFP to the lysosome and/or a portion of TfR undergoing normal lysosomal turnover bringing CYpHER with it.

**Fig. 3.**
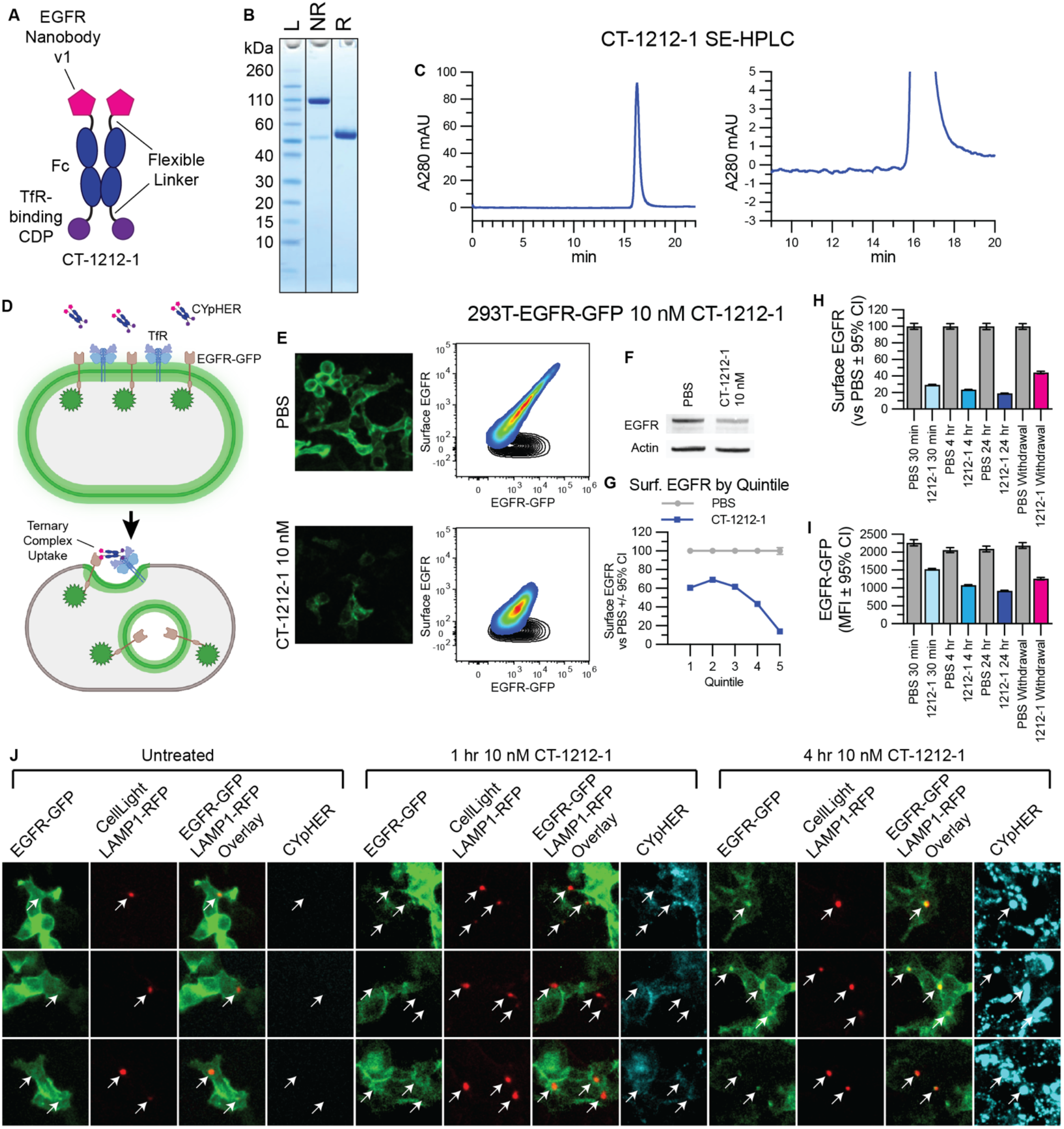
EGFR CYpHER based on VHH nanobody. (**A**) CT-1212-1 design. (**B**) CT-1212-1 SDS-PAGE Coomassie stain. NR: non-reduced. R: DTT-reduced. (**C**) SE-HPLC of CT-1212-1; right is zoomed. (**D**) EGFR-GFP trafficking by CYpHER. (**E**) 293T-EGFR-GFP cells treated 24 hr with PBS or 10 nM CT-1212-1 before either GFP microscopy (left) or flow cytometry after staining for surface EGFR (right). Black contour: unstained cells. (**F**) 293T-EGFR-GFP cells treated with PBS or 10 nM CT-1212-1 for 24 hr and flow sorted for viable (DAPI-) cells prior to Western blotting. (**G**) Same cells and treatment as (**E**), stratified by surface EGFR quintile and normalized. (**H** and **I**) 293T-EGFR-GFP cells dosed with PBS or 10 nM CT-1212-1 for 30 min, 4 hr, 24 hr, or 24 hr followed by 24 hr without drug (“Withdrawal”) were flow analyzed for surface EGFR (**H**) or total EGFR-GFP (**I**) as in (**E**). (**J**) 293T-EGFR-GFP Cells (24 well dish, 500 μL media per well) were treated with 50 μL CellLight Lysosomes-RFP (delivering gene for LAMP1-RFP) for 24 hr, after which they were untreated or treated with 10 nM DyLight 755-labeled CT-1212-1 for 1 or 4 hours and then imaged on the GFP, RFP, and near IR channels. Arrows indicate location of LAMP1-RFP foci (i.e., lysosomes).

For the remaining experiments, we focus most of our protein trafficking data on surface clearance instead of total protein elimination. First, removal of EGFR from the plasma membrane separates it from access to both ligand and downstream signaling modulators like KRas. Second, after endosomal release, actual elimination vs recycling of protein is highly context-dependent, involving cell-specific recycling kinetics and saturable trafficking modulators *(57)*. Third, proteins in the process of synthesis can be detected as total protein but can neither signal effectively nor be accessed by CYpHER.

We explored surface EGFR clearance rates on a panel of non-small cell lung cancer (NSCLC) cell lines which include various drug-resistance mechanisms that can occur in patients: H1650 (EGFR with exon 19 deletion and PTEN knockout), H1975 (EGFR with L858R and T790M mutations [the latter rendering it resistant to 1^st^ generation EGFR TKIs] along with activating G118D mutation in PIK3CA), A549 (wild type EGFR with activated KRas G12S), and H358 (wild type EGFR with activated KRas G12C). They also represent a range of total surface levels and ratios of EGFR and TfR (the latter measured via non-competitive clone OKT9) *(58)* (Fig. 4A). All four lines responded to CYpHER for surface EGFR clearance (Fig. 4B), with rapid kinetics (1 hr) and to a much greater degree than seen with cetuximab, a molecule known to induce surface clearance and overall protein reduction via induction of ubiquitin-mediated uptake *(2, 3)*. We also confirmed that CYpHER remains detectible on the cell surface after media exchange (despite continued cell division and natural TfR turnover), where cetuximab does not (Fig. 4C).

**Fig. 4.**
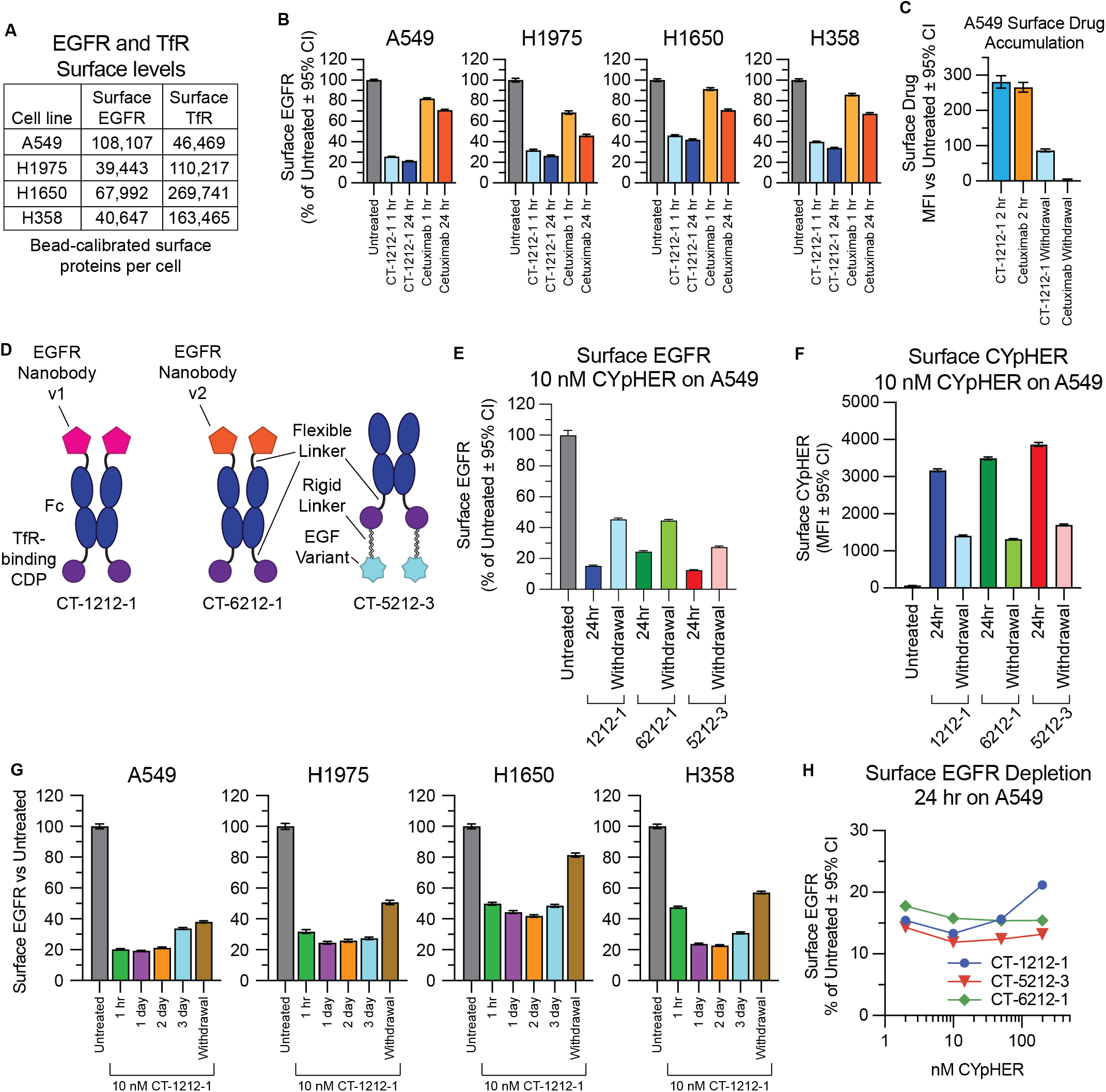
Performance comparison of different EGFR CYpHER designs. (**A**) A549, H1975, H1650, and H358 cells were flow analyzed alongside calibration beads to quantitate surface EGFR and TfR protein levels. (**B**) Normalized surface EGFR levels in the four lines after 1 or 24 hr treatment with 10 nM CT-1212-1 or cetuximab. (**C**) A549 cells incubated with 10 nM CT-1212-1 or cetuximab for 2 hr or for 2 hr followed by 24 hr without drug (“Withdrawal”) followed by staining for human IgG to quantitate surface drug levels. (**D**) EGFR CYpHER designs. (**E**) Surface EGFR levels in A549 cells incubated with 10 nM CYpHER for 24 hr or for 24 hr followed by 24 hr without CYpHER (“Withdrawal”). (**F**) Same treatment as (**E**) but staining for human Fc to quantitate surface CYpHER levels. (**G**) A549, H1975, H1650, and H358 cells untreated or treated with 10 nM CT-1212-1 for 1 hr, 1 day, 2 days, 3 days, or 1 day followed by 1 day without drug (“Withdrawal”) and then analyzed by flow cytometry for surface EGFR levels. (**H**) A549 cells treated for 24 hr with 2 nM, 10 nM, 50 nM, or 200 nM CYpHER and then analyzed by flow cytometry for surface EGFR levels.

### CYpHERs with any of the three engineered EGFR binders clear surface EGFR

The three engineered EGFR binders were incorporated into CYpHER molecules (Fig. 4D). For all three test molecules, the same Fc with a high-affinity TfR-binding CDP (separated by a flexible Gly-Ser linker) was the starting point. The nanobodies were fused via a flexible Gly-Ser linker to the N-terminus of the Fc domain as was done in CT-4212-1; the nanobody’s N-terminus is at the EGFR Domain III interface *(47)*, so this format is optimal for EGFR binding. As described above, using EGFR Nanobody v1 yielded CT-1212-1, while EGFR Nanobody v2 was incorporated into CT-6212-1. Conversely, the C-terminus of EGF is adjacent to the EGFR Domain III interface *(49)*, so fusion via its N-terminus is optimal; it was incorporated in a similar format to that of CT-4212-3, producing CT-5212-3.

Surface EGFR levels in A549 NSCLC cells were reduced after 24 hr CYpHER treatment with all 3 designs, including after media exchange and growth without drug for another 24 hr (“Withdrawal”) (Fig. 4E); as EGFR turnover in the absence of ligand is fairly rapid (∼6-10 hours on most cell lines) *(59, 60)*, this suggested retention of activity via catalytic mechanism. As was seen with CT-1212-1, CYpHER molecules were still present on the surface after a 24 hour drug withdrawal (Fig. 4F).

CT-1212-1 was tested for up to 3 days in the four lung cancer cell lines (A549, H1975, H358, and H1650). All four cancer lines had rapid (within 1 hour) reduction of surface EGFR, ending up with levels between 19% and 45% of untreated after 24 hours (Fig. 4G); this reduction was maintained for at least 3 days. Additionally, 24 hours without drug after 24 hours of treatment (“withdrawal”) still yielded surface EGFR levels markedly lower than untreated cells. We also evaluated surface TfR levels (fig. S4) for potential fluctuation. TfR levels underwent only mild fluctuations that normalized in A549 and H1975 lines. In the lines with the highest TfR levels (H358 and H1650), a more sustained reduction of surface TfR occurred; for both lines, it took 24 hours to reach these low levels, and for both lines, it largely returned to normal after 24 hours of withdrawal. Their higher levels of surface TfR may facilitate CYpHER-induced multimerization, which is known to alter TfR trafficking *(61, 62)*.

We also tested whether we could observe a hook effect. This phenomenon has been documented for some bispecific TPD molecules, wherein target depletion is blunted if the drug concentration is so high that separate drug molecules occupy each respective partner (preventing ternary complex formation) *(63)*. At all doses of CT-1212-1, CT-5212-3, and CT-6212-1 evaluated (2 nM, 10 nM, 50 nM, 200 nM), surface EGFR levels on A549 cells were reduced compared to untreated cells after 24 hours of treatment (Fig. 4H). The degree of EGFR reduction by CT-1212-1 was modestly blunted at 200 nM compared to lower doses, whereas CT-5212-3 and CT-6212-1 show less, if any, variation over this concentration range. This suggests that the effect, if any, is mild and the nature of the molecule (binder and/or modular organization) may have some impact on the balance between ternary complex formation vs separate receptor saturation. Specific effects may also differ by cell line, target, and metabolic state, as it is affected by the relative levels of target and TfR on a given cell.

### CYpHER-driven EGFR intracellular sequestration

We evaluated EGFR trafficking in a knock-in A549-EGFR-GFP cell line using CT-1212-1 and variants thereof (Fig. 5A) that used heterodimeric Fc domains to alter EGFR- and TfR-binding valence. CT-1211-1 has identical EGFR-binding capabilities to CT-1212-1 but only one TfR-binding domain, CT-1112-1 has only one EGFR-binding domain but identical TfR-binding capabilities to CT-1212-1, and CT-1111-1 can only bind one each of EGFR and TfR. All four molecules drove EGFR-GFP from the membrane into intracellular compartments (Fig. 5B), with this internalization activity partially retained after a 24 hour treatment and a 24 hour drug withdrawal. This effect requires both EGFR and TfR binding capabilities, as demonstrated with control molecules containing a control (non-EGFR-binding) nanobody or a control (non-TfR-binding) CDP (Fig. 5C). We confirmed the rapid activity of the mechanism, showing this internalization phenotype after only 20 mins of treatment (Fig. 5D). We also tested holoTF competition, as our TfR-binding CDP binds to the same site on TfR as transferrin *(30)*. Both EGFR-GFP internalization (as seen by microscopy) (Fig. 5E) and surface EGFR clearance (as seen by flow cytometry) (Fig. 5F) were retained in the presence of human holoTF, with only mild suppression of activity at 1000x molar levels of holoTF versus CYpHER. Levels of holoTF in the tumor parenchyma have previously been estimated (as evaluated by quantitative mass spectrometry and ELISA) to be ∼2 nM *(64–66)*, so activity in the tumor microenvironment should not be affected by transferrin.

**Fig. 5.**
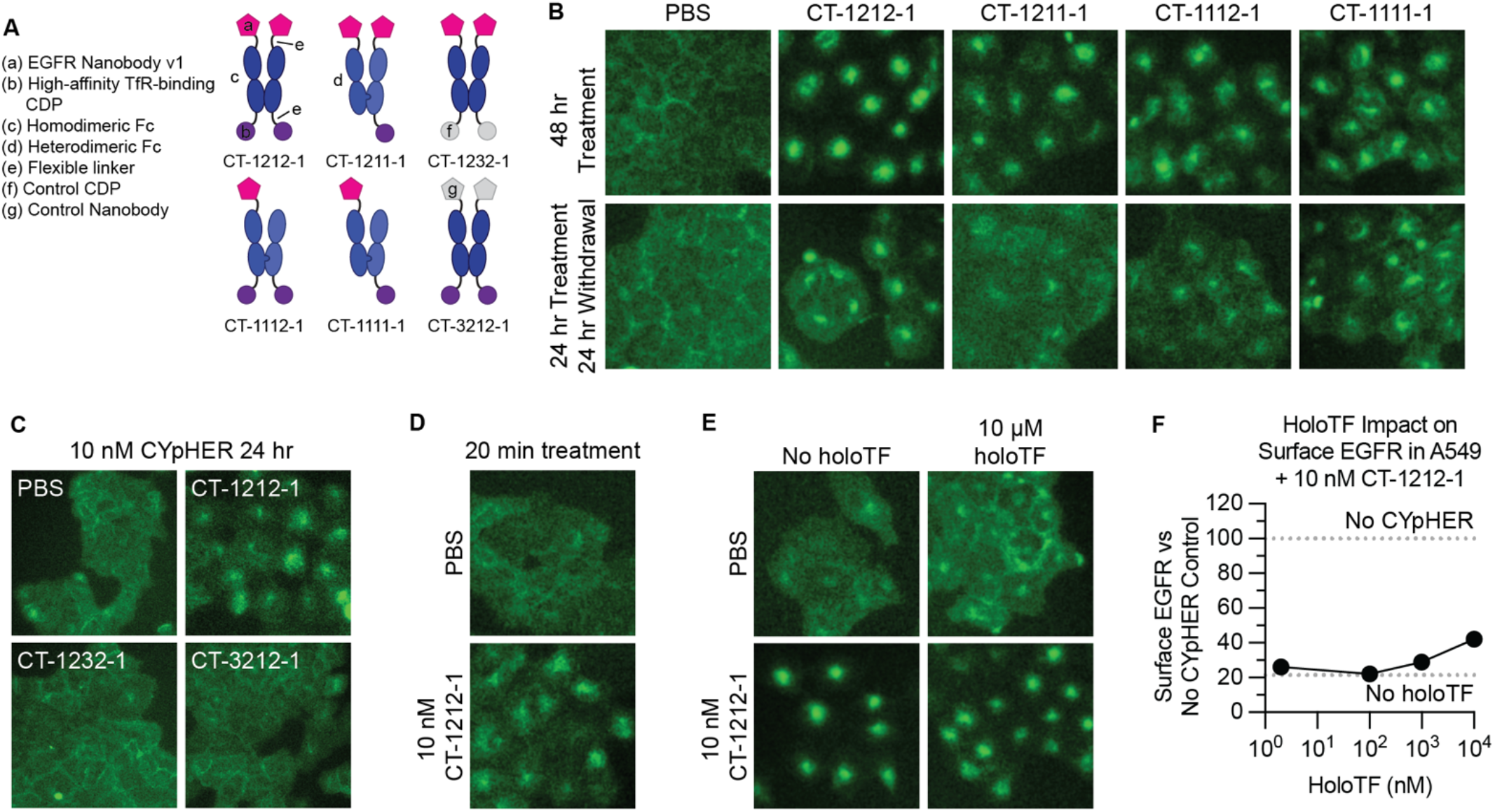
EGFR trafficking upon CYpHER treatment. (**A**) Designs of various EGFR CYpHERs and control molecules. (**B**) A549-EGFR-GFP (knockin) cells treated with PBS or 10 nM CYpHER for either 48 hr with drug, or for 24 hr with followed by 24 hr without drug (“Withdrawal”) and imaged for GFP localization. (**C**) A549-EGFR-GFP cells imaged for GFP localization after 24 hr treatment with 10 nM CYpHER (CT-1212-1) or control molecule (CT-1232-1 or CT-3212-1). (**D**) A549-EGFR-GFP cells imaged for GFP localization after 20 min treatment with PBS or 10 nM CT-1212-1. (**E**) A549-EGFR-GFP cells treated without or with 10 μM human holoTF for 15 mins followed by addition of PBS or 10 nM CT-1212-1 for 4 hr and imaged for GFP localization. (**F**) Same experimental design as in (**E**) except altered amount of holoTF and analyzed by flow cytometry for surface EGFR. Dashed lines indicate quantitation of surface EGFR in untreated cells (upper) or cells treated with 10 nM CT-1212-1 but no holoTF (lower).

### CYpHER catalytic target uptake

Our experiments with drug withdrawal strongly suggest a catalytic mechanism of action where one drug molecule can induce uptake of multiple target molecules, as we see both retention of activity and retention of CYpHER molecules on the surface of cells after 24 hr drug withdrawal. We developed an orthogonal assay system to further evaluate catalytic target uptake (Fig. 6A and fig. S5). In this system, CYpHER facilitates uptake of soluble cargo. We first demonstrated that cells specifically take up soluble target-CYpHER complexes. Cells were exposed to target-saturated CYpHER for 2 hrs, permitting time for CYpHER to drive uptake of soluble target and for CYpHER (via TfR) to cycle in and out of the cell several times. We saw that CYpHER can indeed drive specific uptake of soluble target under these conditions (fig. S5, bars 1 vs 2 [2 hr] and bars 3 vs 5 [24 hr]). Next, the assay was modified to look for uptake of newly introduced target after the initial 2 h uptake followed by removal of all soluble CYpHER from the system. In step 1 of this assay, cells are exposed to target-saturated CYpHER for 2 hrs, with the soluble target being unlabeled. Then the cells are thoroughly rinsed to remove all soluble molecules. After the rinse, cells are exposed to fluorescent soluble target alone for 24 hrs. Any fluorescent target uptake during these 24 hrs, in excess of that seen by cells that were not pre-exposed to CYpHER, is due to the catalytic activity of CYpHER molecules that have already cycled (and released) the previously introduced non-fluorescent target (fig. S5, bars 3 vs 4). With this assay, using various CYpHER designs (Fig. 6B), we saw catalytic CYpHER-driven uptake of cargo with all CYpHER molecules and cell lines tested. Fluorescent soluble EGFRvIII uptake via CT-1212-1 in four NSCLC cell lines, A549, H1650, H1975, and H358 (Fig. 6C), was seen, with catalytic uptake (normalized to passive uptake without CYpHER, which is likely via pinocytosis) at levels that vary by cell line. The degree of catalytic uptake also changed dependent on the nature of the target-binder (Fig. 6D). Additionally, as demonstrated with the same assay using PD-L1 CYpHERs and soluble PD-L1, the format of the CYpHER (Fig. 6E) had an impact. In all cases, uptake of soluble target in excess of passive uptake was seen, demonstrating a catalytic mechanism of action for CYpHER. It should be noted that, with the exception of PD-L1 and H1650 cells (which do not natively express PD-L1), all of these experiments took place in cells that naturally express some amount of the target on the surface, providing competition for soluble cargo binding and uptake. As such, the rate of catalytic soluble cargo uptake is dampened to some degree from its potential maximum, reducing the absolute values observed.

**Fig. 6.**
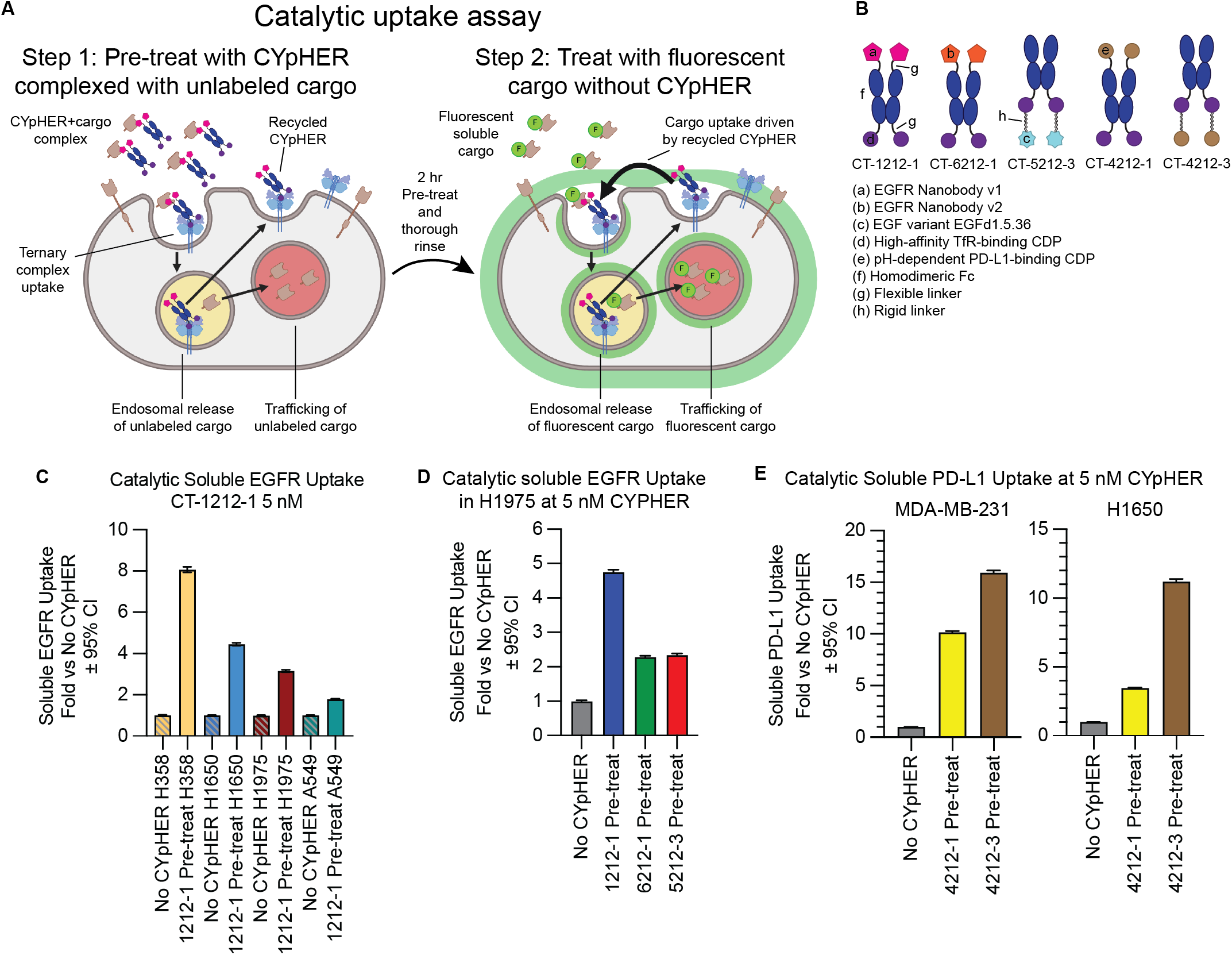
Catalytic soluble cargo uptake by CYpHER. (**A**) Experimental design to quantitate fluorescent soluble cargo uptake in cells pre-treated with cargo-saturated CYpHER. Step 1: CYpHER is saturated with target (2:1 target:binding moiety ratio), then applied to cells for 2 hr. Step 2: After cells are thoroughly rinsed, new fluorescently-labeled soluble target is added to cells. Fluorescence accumulation is increased by CYpHER pre-treatment. (**B**) Designs and elements of CYpHERs used. (**C**) Soluble EGFRvIII uptake after 24 hr incubation with A549, H1650, H1975, or H358 cells either untreated (to quantitate passive uptake) or pre-treated for 2 hr with unlabeled-EGFR-saturated 10 nM CT-1212-1, normalized to each cell line’s untreated uptake levels. (**D**) Soluble EGFRvIII uptake in H1975 cells as in (**C**) comparing CT-1212-1, CT-6212-1, and CT-5212-3. (**E**) Soluble PD-L1 uptake as in (**C**) except with soluble PD-L1 as cargo, comparing CT-4212-1 and CT-4212-3 in MDA-MB-231 (left) or H1650 (right) cells.

### Pharmacodynamic effects of CYpHER *in vitro*

Having established target depletion, we began to investigate how this depletion alters EGFR-mediated signaling and cell growth (Fig. 7). Using various CYpHER and control molecules (Fig. 7A), we investigated ligand-induced signaling, where the level of surface clearance as well as the competitive binding of both our nanobody and EGF variant could have impacts. Stimulating CYpHER-treated (24 hr at 10 nM) cells with 50 ng/mL EGF produced no increased EGFR phosphorylation in CT-1212-1-treated cells (Fig. 7B), suggesting any remaining EGFR on the surface is blocked by the drug or is otherwise signaling-incapable. The valence-altering variants (CT-1211-1, CT-1112-1, CT-1111-1) had the same effect, but control molecules CT-1232-1 (with a non-TfR-binding CDP in place of the high affinity TfR binder) and CT-3212-1 (with a non-EGFR-binding nanobody in place of EGFR Nanobody v1) did not, demonstrating that both TfR- and EGFR-binding functions are necessary together to block ligand-induced activation of EGFR. CYpHER containing the EGF variant was also tested (Fig. 7C); in contrast to the nanobody-containing CYpHER series, CT-5212-3-treated cells showed some residual EGFR phosphorylation in response to EGF, suggesting the EGF variant on the CYpHER provides less complete blockade of the EGF-binding site of the residual surface-resident EGFR. Further affinity maturation of the variant may improve its ability to competitively inhibit EGF stimulation under these conditions. At the same time, we saw that the EGF variant itself (in the form of Fc fusion CT-5200-6) did not induce EGFR phosphorylation, confirming its capacity as a dominant-negative EGF.

**Fig. 7.**
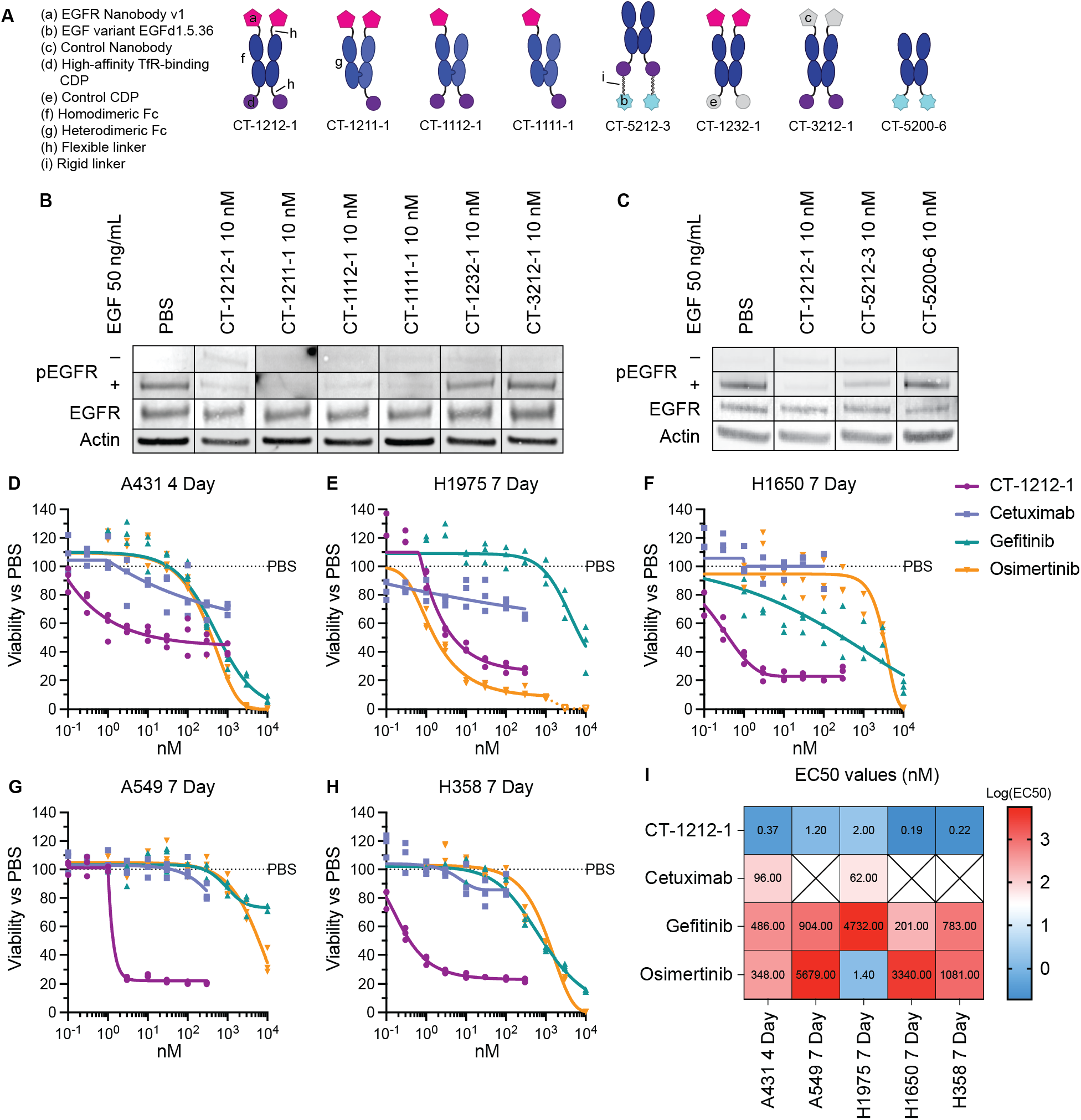
Pharmacodynamic effects of CYpHER. (**A**) Designs of various EGFR CYpHERs and control molecules. (**B** and **C**) A549 cells (unsorted, thus including live and dead cells) treated for 24 hr with PBS, 10 nM CYpHER, or 10 nM control molecule followed by no treatment (“–”, EGFR, Actin) or addition of 50 ng/mL EGF for 30 min (“+”) and analyzed by Western blot for pY1068 EGFR (“–” and “+”), total EGFR, or actin. (**D** to **H**) Triplicate 96 well plate growth for 4 days (A431) or 7 days (all others) with single dose (no media exchange) of CT-1212-1, cetuximab, gefitinib, or osimertinib in A431 (**D**), H1975 (**E**), H1650 (**F**), A549 (**G**), and H358 (**H**) cells. After treatment, cell levels per well were quantitated by CellTiter-Glo 2.0 assay. (**I**) EC50 values of the experiments in (**D** to **H**) from asymmetric sigmoidal (5PL) curve fit. Empty “X” box indicates no effect, as defined by failure to suppress growth by 20% at any dose tested.

As head and neck cancer (e.g., HNSCC) and lung cancer (e.g., NSCLC) are commonly treated with EGFR-targeting agents *(6, 67)*, CT-1212-1 was tested for the ability to suppress cell growth in several relevant cell lines: head and neck cancer line A431 (with a massive duplication of the EGFR gene), and the four NSCLC lines H1650, H1975, A549, and H358. All of these lines were tested for growth suppression dose response alongside clinical EGFR-targeted drugs cetuximab (EGFR mAb), gefitinib (1^st^ generation EGFR TKI), and osimertinib (3^rd^ generation EGFR TKI) (Fig. 7, D to I). CT-1212-1 suppressed growth across all cancer types and mutational profiles tested. Moreover, It had higher potency (lower growth suppression EC50 values) than the three clinical comparators in all five lines, with the exception of comparable potency with osimertinib in the H1975 cell line that carries the T790M mutation against which osimertinib was originally developed *(68)*.

CT-1212-1 strongly suppressed cell growth but did not result in complete cell killing *in vitro*. Many targeted therapies, including osimertinib, can be cytostatic *in vitro* but cause tumor regression *in vivo (69)*. In a complete biological system *in vivo*, EGFR disruption upregulates pro-inflammatory stress pathways, leading to inflammatory cytokine secretion, immune cell infiltration, and reduction of regulatory T cells, all of which support an anti-tumor response *(70)*. Likewise, TKIs can show cytostatic activity *in vitro* at doses that are selective for their target (as seen in osimertinib in H1975 cells at lower doses [Fig. 7E]), but at high doses, the TKIs disrupt so many off-target kinases that cell viability is broadly impacted.

As many EGFR-targeted therapeutics (mAbs and TKIs) demonstrate skin toxicities *(7, 71)* due to the sensitivity of keratinocytes to EGFR suppression *(72)*, the activity of CT-1212-1 was also tested in primary human dermal keratinocytes (fig. S6). First, we examined levels of both TfR and EGFR on these cells. TfR levels were much lower on these normal cells than on the cancer lines we tested, but EGFR levels are quite high on this sensitive cell population (fig. S6A). Surface EGFR clearance was observed with CT-1212-1, but with much slower kinetics than seen in the cancer lines (likely due to much lower levels of surface TfR), and no reduction in surface TfR levels was seen (fig. S6B). In primary keratinocyte growth inhibition assays (fig. S6C), CT-1212-1 demonstrated similar properties to cetuximab, and the TKIs had similar profiles as their activity on the cancer cell lines. In cancer patients, the therapeutic window is the drug concentration range where efficacy against cancer is seen while also avoiding unacceptable toxicity. Knowing that skin toxicity is a prominent concern for EGFR targeted therapeutics *(73)*, we compared potency at suppressing cancer cell growth to keratinocyte growth suppression (fig. S6D). The potency against cancer vs the potency against keratinocytes was higher for CT-1212-1 than for any of the three clinical drugs in the five cell lines, with the exception of osimertinib in the EGFR T790M-carrying H1975 line.

### *In vivo* CYpHER pharmacokinetics and pharmacodynamics

The CYpHER candidates were designed as Fc fusions, with advantages including extending *in vivo* pharmacokinetic (PK) properties as well as ease in manipulating avidity (monovalent vs bivalent binding to either target or TfR). To investigate the former, we dosed NCr nu/nu mice with 1.5 mg/kg CT-1212-1, CT-1222-1, CT-1211-1, and CT-1232-1 (Fig. 8). The nanobody does not cross-react with murine EGFR, so only the murine cross-reactive TfR-binding CDP would be expected to influence PK apart from the Fc domain. CT-1212-1 has two high-affinity TfR-binding CDPs, CT-1222-1 has two medium-affinity TfR-binding CDPs, CT-1211-1 has one high-affinity TfR-binding CDP, and CT-1232-1 has no TfR-binding capability (Fig. 8A). As measured by ELISA (Fig. 8B), serum levels of the CYpHER molecules demonstrated serum half-lives between 41 and 88 hours at this dose, with the longest belonging to the non-TfR-binding molecule CT-1232-1. It is likely that TfR binding is increasing clearance, a phenomenon seen in other studies *(61, 74)*, even with the Fc otherwise extending serum residence. Considering that CYpHERs exhibited potency on cancer cells at concentrations as low as 0.2 nM and EGFR surface clearance at concentrations as high as 200 nM, the data indicates that serum levels in a therapeutic range may be readily attained with infrequent dosing. The biodistribution to the target tissue (e.g., tumor), as well as the durability of activity given catalytic target clearance in cells even when CYpHER is removed from extracellular fluid, are still to be investigated.

**Fig. 8.**
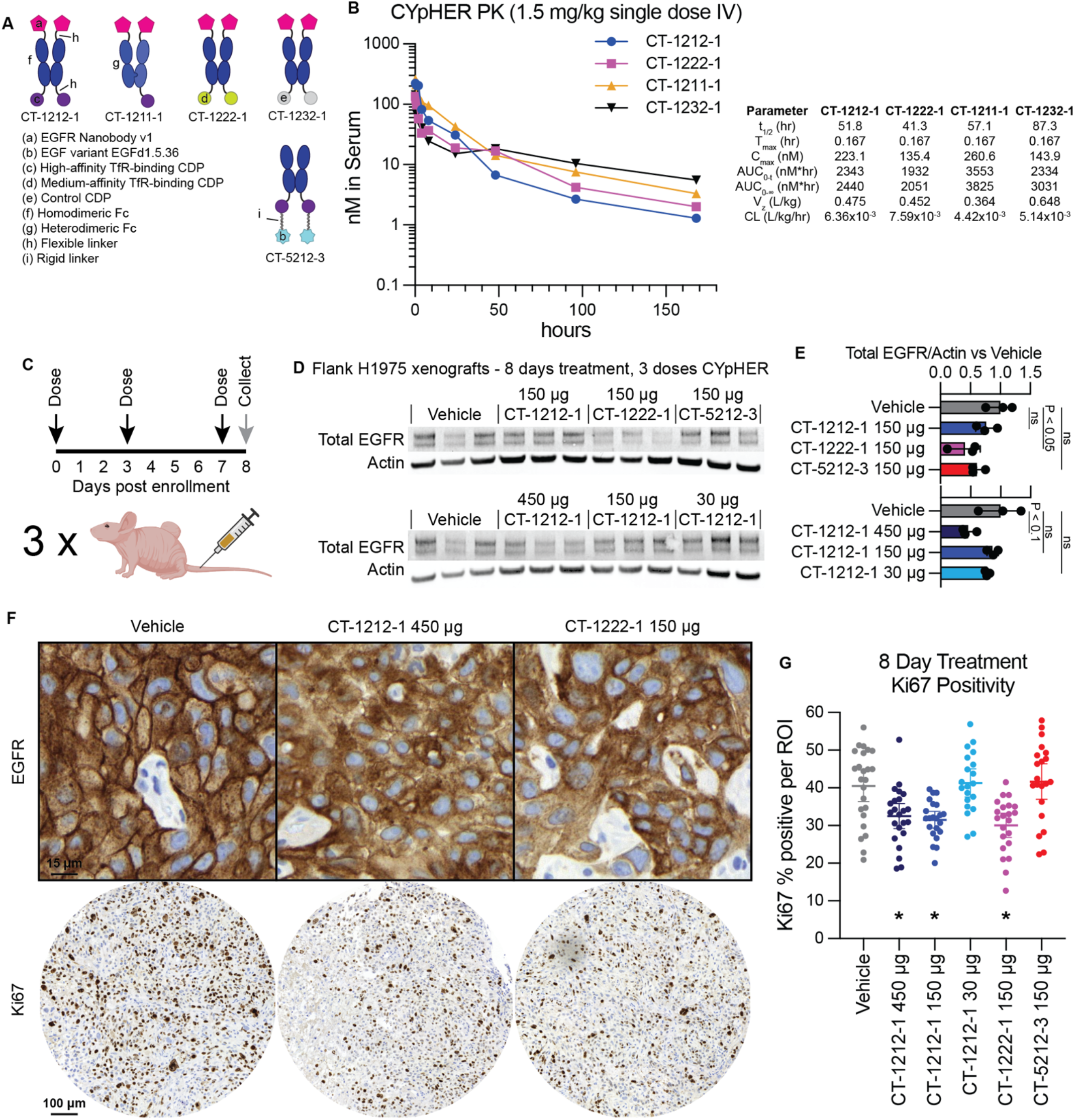
Pharmacokinetics and pharmacodynamics of CYpHER in mice. (**A**) Designs of various EGFR CYpHERs and control molecules. (**B**) NCr nu/nu mice were dosed with 1.5 mg/kg CT-1212-1, CT-1211-1, CT-1222-1, or CT-1232-1 IV. Serum samples were taken after 10 min, 30 min, 1 hr, 2 hr, 4 hr, 8 hr, 24 hr, 48 hr, 96 hr, or 168 hr. Three mice per time point were analyzed. Serum samples were quantitated by ELISA for human Fc domain in technical triplicate. Molecules exhibited a normal biphasic distribution curve, and as such, PK parameters were determined by non-compartmental analysis for IV bolus dosing using PKSolver 2.0. (**C**) Experimental design for tumor implantation and dosing. Female athymic nude mice (Foxn1^nu^) were implanted (subcutaneous flank) with 5×10^6^ H1975 cells. After 21 days, mice were enrolled and dosed IV on days 0 (enrollment day), 3, and 7. On day 8, tumors from three mice per dosage group were harvested and split in half for Western blot lysis or for histology. (**D**) Western blot analysis of total EGFR and actin. (**E**) Quantitation of the blots from (**D**). (**F**) IHC (hematoxylin/DAB) for total EGFR (top) and Ki67 (bottom) in vehicle, CT-1212-1 450 μg, and CT-1222-1 150 μg groups. Full EGFR fields can be found in fig. S7. (**G**) Quantitation of Ki67 positivity, derived from 6-9 regions of interest (ROI) per tumor, three tumors per group, pooled for analysis. *: P < 0.01 vs vehicle.

We next tested mice implanted with flank H1975 xenografts and treated for 8 days with CYpHER for any observed pharmacodynamic (PD) effects (Fig. 8C). Female athymic nude mice were implanted with 5×10^6^ H1975 cells. After 21 days of growth, mice received three doses of CYpHER on days 0 (enrollment day), 3, and 7. CYpHERs and doses administered were: CT-1212-1 450 μg/dose; CT-1212-1 150 μg/dose; CT-1212-1 30 μg/dose; CT-1222-1 150 μg/dose; and CT-5212-3 150 μg/dose. On day 8, tumors were harvested for Western blotting and histology. Western blotting (Fig 8, D and E) demonstrated a reduction (P < 0.05, as normalized to actin) in total EGFR by CT-1222-1 150 μg/dose, and a trend towards EGFR reduction (P < 0.1) by CT-1212-1 450 μg/dose. Histology for EGFR and Ki67 (Fig 8, F and G) demonstrated two phenomena. First, most fields from the CT-1212-1 450 μg/dose and CT-1222-1 150 μg/dose tumors demonstrated altered localization of EGFR (Fig. 8F), visible as marked reduction of membrane EGFR (DAB stain) relative to intracellular levels. Second, automated Ki67 quantitation (Fig. 8G) demonstrated that 8 days of treatment caused a reduction (P < 0.01) in the proliferation marker Ki67 (percent Ki67 positivity) in the CT-1212-1 450 μg/dose, CT-1212-1 150 μg/dose, and CT-1222-1 150 μg/dose groups vs vehicle.

## Discussion

Catalytic TPD via CYpHER is a novel approach to durably depleting disease-driving proteins, capable of altering target trafficking and cell behavior both *in vitro* and *in vivo*. CYpHER leverages TfR, a protein that has well-characterized and rapid recycling kinetics, generalizing the activity across many tumor and cell types. Furthermore, TfR is overexpressed and required for cancer cell growth, increasing potential tumor accumulation and reducing the risk of acquired drug resistance. TfR also delivers cargo across the blood-brain barrier, adding CNS proteins as potential targets. Through engineering the target-binding end for pH-dependent release, CYpHER is not reliant on other enzymes for function, and its catalytic activity permits one drug molecule to clear multiple target proteins with more durable function. Lastly, the drug molecules are proteins produced by standard recombinant expression, requiring no chemical modifications and using binder modalities found in other clinically-approved molecules.

CYpHER is a promising approach for the inexorable challenge of EGFR-driven cancer for many of the reasons described for RTKs in general. Established therapeutics, particularly TKIs, can effectively induce tumor regression, but nearly all patients experience relapse with resistance mutations, often within months *(6)*. EGFR-targeting CYpHER could simultaneously avoid many of these common resistance mechanisms and could offer targeted therapy to patients who have developed resistance to frontline treatment *(75)*. Our data on relevant mutations supports this possibility. Point mutations to disrupt drug activity are straightforward and commonplace in TKI treatment (T790M in 1^st^ generation TKIs like gefitinib, C797S in 3^rd^ generation TKIs like osimertinib) *(7, 8)*, but CYpHER should be able to deplete any of these variants. Tumor upregulation of EGFR is commonly seen in response to targeted therapeutic treatment, but in cell populations with varying target expression levels, we’ve seen CYpHER have stronger surface and overall target elimination effects in cells that have higher target expression, so clones with higher EGFR expression are less likely to lead to relapse. Meanwhile, upregulation of other RTKs that can cross-phosphorylate EGFR are also less likely to succeed with a CYpHER approach, as elimination of EGFR from the surface removes its use as both kinase and substrate for these heterodimer partners. ECD mutations to avoid drug binding are less common in response to biologic drugs against EGFR *(76, 77)*, likely because of the greater interface surface area that often requires disruption of multiple points of interaction to meaningfully reduce binding. CYpHER, as a platform, can approach numerous difficult targets in oncology and CNS disease. The ErbB family of RTKs (e.g., EGFR, HER2, ERBB3) are all associated with driving cancer and/or inflammation in various tissues and settings, alongside other growth factor and cytokine receptors (e.g., MET, FGF receptors, IGF-1 receptors, interleukin receptors). All feature multiple functions, which can include homodimerization, heteroassociation, kinase function, and/or scaffolding for signal transduction. Any of these functions can be approached by conventional therapeutics, but protein elimination is the only way to completely neutralize all possible disease-causing functions. Knockdown and knockout strategies are advancing in the clinic, but TPD approaches like CYpHER can be effective with less pharmacological complexity. Additionally, access to the CNS via TfR-mediated transcytosis is an exciting avenue for dealing with difficult tasks like clearing neurodegeneration-associated misfolded proteins (e.g., amyloid, tau, huntingtin) or their inflammatory mediators. Metastasis to the CNS is also a common cause for cancer progression on otherwise effective therapy, and TfR-mediated CNS access may prevent this mechanism of recurrence.

In conclusion, CYpHER adds a powerful entry to the TPD field. With catalytic functionality, broad target applicability, good assembly and production, potent and durable alteration of target trafficking, and demonstrable *in vivo* activity, proteins for which traditional targeted therapeutics have struggled may be approachable, with promising implications to our most insidious and intransigent diseases.

## Methods

### Recombinant proteins, antibodies, and co-stains/secondary antibodies

Recombinant proteins used for surface display flow cytometry and for catalytic uptake of soluble proteins were as follows: biotinylated His-Avi-tagged human PD-L1 ectodomain (ACROBiosystems PD1-H82E5); biotinylated His-Avi-tagged human full-length EGFR ectodomain (VHH nanobody analysis, ACROBiosystems EGR-H82E3); biotinylated His-Avi-tagged human EGFRvIII ectodomain (EGF variant analysis and soluble EGFR uptake, ACROBiosystems EGR-H82E0); biotinylated His-Avi-tagged human TfR (TfR cross-reactivity, ACROBiosystems TFR-H82E5); His-tagged mouse TfR (TfR cross-reactivity, R&D Systems 9706-TR-050; note that this was produced in Chinese hamster ovary [CHO] cells, as opposed to all other recombinant proteins that were produced in human HEK cells).

Primary antibodies for surface protein staining were as follows: EGFR, clone 199.12 (ThermoFisher MA5-13319); TfR, clone OKT9, APC-labeled (ThermoFisher 17-0719-42); PD-L1, clone 22C3 (Agilent M365329-1). Primary antibodies for Western blotting are as follows: rabbit anti-EGFR (Cell Signaling Technology 2646); rabbit anti-phospho-Y1068 EGFR (Cell Signaling Technology 3777); goat anti-actin (Abcam ab8229). Secondary antibodies or co-stains were as follows: Alexa Fluor 647-conjugated streptavidin (surface display staining other than TfR human/mouse cross-reactivity, ThermoFisher S21374); iFluor 647-conjugated anti-His-tag antibody (pilot CYpHER detection and TfR human/mouse cross-reactivity, Genscript A01802); Alexa Fluor 647-conjugated anti-Fc Fab (Surface protein and CYpHER quantitation, ThermoFisher Zenon Labeling Kits, mouse IgG2a for anti-EGFR 199.12 quantitation [ThermoFisher Z25108], mouse IgG1 for anti-PD-L1 22C3 quantitation [ThermoFisher Z25008], human IgG for CYpHER quantitation [ThermoFisher Z25408]); iFluor 647-conjugated sAvPhire monovalent streptavidin (catalytic soluble protein uptake, Millipore Sigma SAE178-100UG); iFluor 488-conjugated sAvPhire monovalent streptavidin (catalytic soluble protein uptake, Millipore Sigma SAE176-100UG); IRDye 680RD Donkey anti-goat (Western blotting, LI-COR 926-68074); IRDye 800CW Donkey anti-rabbit (Western Blotting, LI-COR 926-32213).

### EGF variant design using Rosetta

A co-crystal structure containing EGF and EGFR (PDB 1IVO) was processed to separate EGF (chain C) and EGFR domain III (chain A, residues 311-510). They were combined into a single PDB, which was used as the input for Rosettascripts *(53)* using proprietary XML scripts optimized for CDP redesign. 1000 unique variants were designed and scored using an interface analysis script, with 488 that had favorable scoring parameters incorporated into a mammalian surface display screening library.

### Mammalian surface display

293F cells (ThermoFisher R79007) were grown in FreeStyle 293 expression medium (ThermoFisher 12338018) in 37°C, 8% CO_2_ humidified shaking incubators. Proteins were surface displayed via transient transfection (singleton testing) or lentiviral transduction (pooled screening) using vector SDGF *(46)*, in which displayed proteins have a free C-terminus, or a variant thereof where the displayed protein has a free N-terminus and is connected to C-terminal GFP by a Type 1 transmembrane domain derived from human CD28. The parental C-terminal display vector was used for experiments involving CDPs (including EGF and variants thereof), while the N-terminal display variant was used for experiments involving VHH nanobodies. General growth, transfection, staining, sorting, and data interpretation methods were previously published *(30, 45, 46)*. Staining either took place with monovalent (TfR-binding CDP or PD-L1-binding CDP work) or tetravalent (VHH nanobody or EGF variant work) protocols, with binder concentrations varying depending on the assay: 100 nM for diversity library screening (Primary EGF Rosetta variant library), 20-100 nM for maturation (EGFd1.5 affinity maturation, VHH nanobody His-doped variant library, PD-L1 binder pH maturation), and 10-50 nM for singleton validation stains. Testing for pH-dependent release involved the conventional staining protocols, but after target protein incubation, cells were pelleted and resuspended in cold pH 7.4 PBS or pH 5.5 citrate-phosphate saline buffer for 5 mins, followed by pelleting at 500xg for 5 mins (combined 10 mins incubation). Cells were then resuspended in buffer for the next step (fluorescent co-stain for monovalent staining protocols, Flow Buffer [PBS with 0.5% bovine serum albumin and 2 mM EDTA] for tetravalent staining protocols). Flow cytometry took place on Becton Dickson FACSAria III or on Sony SH800S instrumentation.

### Recombinant protein production and analysis

Pilot CYpHER (fig. S1) molecule was produced as previously described *(33, 78)*. All other CYpHER molecules were produced by transient expression in suspension HEK293 cells (ThermoFisher GeneArt) and purified either via Ni-NTA pull-down as previously described *(33, 78)* or by Protein A columns (Cytiva 28985254 [pre-packed Protein A columns] and 28903059 [buffer kit]) as per manufacturer’s protocol. Proteins were buffer exchanged (Sephadex G25 desalting columns, Cytiva 17085101) into PBS and aliquoted for storage at -80°C. SDS-PAGE (4-12% Bis-Tris 1 mm thickness, ThermoFisher NP0321BOX or NP0323BOX) was run with MES buffer (ThermoFisher NP0002) at 180V for 50 min prior to Coomassie stain. SE-HPLC was performed on Agilent instrumentation using a TSKgel G3000SWXL column (Tosoh 08541). Mobile phase was 50 mM acetate pH 5.0, 100 mM NaCl, 100 mM arginine, 5% EtOH. Flow rate for the run was 0.5 mL/min. 100 μg protein was loaded.

### Surface Plasmon Resonance (SPR) Interaction Analyses

SPR experiments were performed at 25°C on a Biacore 3000 instrument (Cytiva) with a CM3 sensor chip and 10 mM Hepes, pH 7.4 or 5.8, 150 mM NaCl, 3 mM EDTA, 0.05% Surfactant P20, and 0.1 mg/mL BSA as the running buffer. Goat anti-human IgG capture antibody (Jackson ImmunoResearch Laboratories, Inc., West Grove, PA; 109-005-098) was immobilized to flow cells of the sensor chip using standard amine coupling chemistry to a level of 3,800 RU to 3,900 RU. Fc fusion molecules were captured to a level of 70 RU (pH-dependent release VHH) and 55 RU (high affinity at low pH VHH). EGFR concentration ranges were 150 nM to 1.85 nM (pH-dependent release VHH Fc fusion) or 50 nM to 0.617 nM (high affinity at low pH VHH Fc fusion). Each analyte concentration series was run in duplicate and in mixed order, as a means of assessing the reproducibility of binding and managing potential systematic bias to the order of injection. Multiple blank (buffer) injections were run and used to assess and subtract system artifacts. The association phases were monitored for 240 s and the dissociation phases were collected for 600 s, at a flow rate of 30 μL/min. The surface was regenerated with 10 mM glycine, pH 1.5 for 30 s, at a flow rate of 30 μL/min. The data were aligned, double referenced, and fit using Scrubber v2.0 software (BioLogic Software Pty Ltd, Campbell, Australia), which is an SPR data processing and non-linear least squares regression fitting program. The 240 s association phase data and the 600 s dissociation phase data were globally fit to the 1:1 binding model, to determine the kinetic parameters.

### Cancer cell line and primary keratinocyte surface protein flow cytometry

Cell lines were grown by conventional adherent cell culture in 37°C, 5% CO_2_ humidified incubators using DMEM + 10% FBS and 1x antibiotic/antimycotic (293T-EGFR-GFP, MDA-MB-231-PD-L1-GFP, A431) or RPMI + 10% FBS and 1x antibiotic/antimycotic (A549, H1975, H1650, H1650-PD-L1-GFP, H358). Cells were lifted by removing media, rinsing with room temperature PBS, and incubating with TrypLE Express (ThermoFisher 12605036) until detachment. For staining, following detachment, enzyme was inhibited with complete culture medium, and cells were pelleted at 500xg for 5 min. Cells were resuspended in cold Flow Buffer; for surface EGFR or TfR quantitation, the buffer contained either 10 nM primary antibody (either conjugated with Alexa Fluor 647 [anti-TfR] or pre-labeled with Zenon labeling kit according to manufacturer’s instructions [anti-EGFR]) or an equivalent volume of a mock labeling reaction (using Zenon labeling kit reagents but with flow buffer in place of primary antibody); for surface CYpHER detection, Zenon human IgG detection reagent was added as if it were a primary antibody to 10 nM. Cells were incubated in this staining solution on ice for 30 mins, pelleted at 500xg for 5 mins, and resuspended in fresh, cold Flow Buffer containing 1 μg/mL DAPI immediately prior to flow analysis.

Quantitation of protein copies per cell used this same staining protocol, but staining was done alongside Simply Cellular anti-mouse IgG microspheres (Bangs Laboratories 810), with the lot comprised of beads with an average capacity of 73,000 antibodies. For surface quantitation, fluorescence levels of stained cells were compared to that of the IgG microspheres stained in parallel (subtracting values of cells or beads stained without primary antibody), multiplying the cell lines’ average fold-difference vs the IgG microsphere fluorescence by 73,000 to arrive at the protein copies per cell.

### Catalytic soluble protein uptake

Cells were grown in 24 well plates at 500 μL media per well. For step 1 (CYpHER pre-treatment), cell culture media containing 200 nM recombinant biotinylated target protein ectodomain (EGFRvIII or PD-L1) and 200 nM iFluor 488-conjugated sAvPhire monovalent streptavidin (only 647 channel was analyzed, so this is considered an “unlabeled” stain but used to ensure equivalent target behavior) with or without 50 nM CYpHER were prepared at least 30 min before use, enough for >50 μL per well. 50 μL of this solution was then added to cells growing in 450 μL fresh media, with final concentrations of 20 nM total target protein/streptavidin complex and 5 nM target/streptavidin-saturated binding moieties on CYpHER molecules. Under these conditions, for CYpHER-inclusive conditions, CYpHER molecules saturated with target protein bind to cells and begin cycling through endosomes, trafficking with TfR. After 2 hours, media was removed and cells were gently rinsed twice with PBS before 450 μL fresh media is added. For step 2 (catalytic uptake), cell culture media containing 100 nM recombinant biotinylated target protein ectodomain (EGFRvIII or PD-L1) and 100 nM iFluor 647-conjugated sAvPhire monovalent streptavidin was prepared alongside the step 1 buffers above. 50 μL of this solution was then added to each well for a final concentration of 10 nM target protein/streptavidin complex. Cells were incubated for 24 hrs, during which time cells exposed previously to target/streptavidin-saturated CYpHER that has cycled through the cell and released the target could use the available CYpHER molecules for target binding and uptake. After 24 hrs, cells were rinsed and lifted as above for flow analysis (which included no cell staining, apart from resuspension in Flow Buffer containing 1 μg/mL DAPI). The 647 channel was used for quantitating target uptake.

### Microscopy

Fluorescent cell images were taken on an EVOS M5000 equipped with DAPI (Ex 357/44, Em 447/60), GFP (Ex 470/22, Em 525/50), Texas Red (Ex 585/29, Em 628/32), and Cy7 (Ex 716/40, Em 794/32) cubes. All images within a given experiment were taken with the same light / exposure / gain settings. Image processing took place in ImageJ2 with the Fiji package. When used, background subtraction, contrast enhancement, and gaussian blur filtration (to reduce pixelation) were always done identically between all images within a given experiment using scripts to ensure consistency. CT-1212-1 labeling with DyLight 755 labeling kit (ThermoFisher 84539) was as per manufacturer’s protocol.

### Western blotting

Lysates were prepared with RIPA buffer (ThermoFisher 89900) containing protease/phosphatase inhibitors (Pierce A32959, 1 tablet per 10 mL) and nuclease to reduce viscosity (Pierce 88701, used at 1:1000). Proteins were quantitated by BCA assay (Pierce 23225). SDS-PAGE (4-12% Bis-Tris 1 mm thickness, ThermoFisher NP0321BOX or NP0323BOX) was run with MES buffer (ThermoFisher NP0002) at 180V for 50 min. Gels were transferred to PVDF membrane via iBlot system (ThermoFisher IB401031) on pre-set Program 3. Blotting took place with the LI-COR system (LI-COR 927-66003 [TBS-based blocking and diluent buffers] and ThermoFisher 28360 TBS-Tween 20 wash buffer) for imaging on a LI-COR Odyssey instrument. Primary antibody concentrations were 1:1500 (total EGFR) or 1:3000 (phospho-Y1068 EGFR; actin). Secondary antibody concentrations were 1:15,000. Note: quantitation of bands with bubbles (e.g., Fig. 8D) involved quantitating the unobstructed portion of the band, calibrating against the same portions of the Vehicle bands, and extrapolating.

### Cell viability dose response

Cells were passaged into 96 well plates at 500 or 1000 cells/well, with amounts determined by cell titration to produce final confluence of 30-50% in vehicle treatment. 1 day after plating, wells were dosed with vehicle or compound at indicated concentrations (from 10x stocks prepared separately) in technical triplicate. Cells were grown for 4 days (A431 cells) or 7 days (all others) and then viability assessed by CellTiter-Glo 2.0 per manufacturer’s instructions. Luminescence data were processed in GraphPad Prism v10, normalized to vehicle (and with vehicle used in curve fits as 0.001 nM), and EC50 values are calculated via log-transformed asymmetric sigmoidal (5PL) curve fit with the following deviations from default settings: Max iterations = 10,000, weighing method = weigh by 1/Y^2^, constrain S > 0, constrain Hill Slope < 0, constrain top < 110%, constrain bottom > 0%.

### *In vivo* pharmacokinetic analysis

PK work was performed at Charles River Laboratories. For each test article, 12 female NCr nu/nu mice received single IV doses of 1.5 mg/kg test article in PBS (or only PBS vehicle). In groups of 3, mice were bled at 10 min, 30 min, 1 hr, 2 hr, 4 hr, 8 hr, 24 hr, 48 hr, 96 hr, and 168 hr. 12 mice produced these samples: one trio of mice was bled at 10 min, 4 hr, and 96 hr; one trio was bled at 30 min, 8 hr, and 168 hr; one trio was bled at 1 hr and 24 hr; and one trio was bled at 2 hr and 48 hr. Serum samples were snap-frozen and stored at -80°C until analysis using an in-house ELISA method.

### *In vivo* tumor implantation and dosage

Tumor implantation and dosage was performed at Seattle Children’s Research Institute. All mice were maintained in accordance with the National Institutes of Health Guide for the Care of Laboratory Animals with approval from the Seattle Children’s Research Institute, Institutional Animal Care and Use Committee (protocol ACUC00682). Female Athymic Nude mice (Foxn1^nu^) were purchased from Inotiv Laboratories (#069) and housed under specific pathogen free conditions. NCI-H1975 lung tumor cells were purchased from ATCC (CRL-5908) and verified human pathogen and mycoplasma free. CYpHER proteins CT-1212-1, CT-1222-1, CT-5212-3, and CT-6212-1 were produced as described above, formulated in phosphate-sucrose buffer and confirmed to meet endotoxin specifications. 5×10^6^ tumor cells in PBS were inoculated in the subcutaneous space on the right flank of seven weeks old mice. Study enrollment was done en mass on day 0, 21 days after tumor implantation when the average tumor volume per group was 275 mm^3^. Six mice were randomly assigned to each treatment group, normalizing for equal starting tumor volume. Vehicle or therapeutic were administered as a 200 μL bolus via tail vein injection on days 0, 3, and 7. Tumor volume and body weight were recorded on days 0, 2, 4, and 7. The study ended on day 8. Mice were removed from the study early if ulcerations developed on the tumor surface. All mice were group-housed with unrestricted mobility and free access to food and water for the duration of study.

### Immunohistochemistry

Immunohistochemistry was performed by the Fred Hutchinson Cancer Center Experimental Histopathology shared resource (NIH grant P30 CA015704). Paraffin sections were cut at 4 μm, air dried at room temperature overnight, and baked at 60°C for 1 hr. Slides were stained on a Leica BOND Rx autostainer (Leica, Buffalo Grove, IL) using Leica Bond reagents. Endogenous peroxidase was blocked with 3% hydrogen peroxide for 5 min followed by protein blocking with TCT buffer (0.05M Tris, 0.15M NaCl, 0.25% Casein, 0.1% Tween 20, and 0.05% Proclin 300 at pH 7.6 ± 0.1) for 10 min. Primary antibodies were incubated for 1 hr and polymers were applied for 12 mins, followed by Mixed Refine DAB (Leica DS9800) for 10 min and counterstained with Refine Hematoxylin (Leica DS9800) for 4 min after which slides were dehydrated, cleared and coverslipped with permanent mounting media. IHC antibodies: rabbit anti-EGFR (clone SP84, Cell Marque #14R-15) at 1:25 and mouse anti-Ki67 (clone MIB1, Dako #M7240) at 1:50 with ME Kit, a flexible mouse-on-mouse immunohistochemical staining technique adaptable to biotin-free reagents, immunofluorescence, and multiple antibody staining (https://pubmed.ncbi.nlm.nih.gov/24152994/). Secondary polymer was Power Vision Rabbit HRP Polymer.

### CYpHER ELISA

Goat anti-human Fab’(2) (Jackson ImmunoResearch 109-006-190) was used to coat black Maxisorp plates (Thermo 437111) at 500 ng/mL in 100 μL/well incubated overnight, up to 3 days. After coating, all steps were performed at room temperature with extreme care to ensure all wells of a plate were exposed to the same environmental temperature, avoiding gradients across the plates. Wells were aspirated and blocked with 200 μL PBS containing 3% BSA and 0.1% Tween 20 for 2 hr. After aspiration and three rinses with 250 μL/well PBS containing 0.05% Tween 20, samples and standards were applied. For standards, a ten-point standard curve, covering final in-plate concentrations from 300 ng/mL to 0.015 ng/mL, was prepared for each plate using normal mouse serum as diluent. Each sample and standard were then diluted 1:100 (samples and standards) or 1:1000 (samples only as additional dilution series) into Diluent Buffer (PBS containing 1% BSA and 0.1% Tween 20) prior to addition to the plate and allowed to equilibrate to room temperature. Samples were added to the blocked, rinsed plate (100 μL/well) and incubated for 1 hr. After rinsing as above, wells were then incubated with 100 μL mouse anti-human (clone JDC-10, Abcam ab99760) at 50 ng/mL in Diluent Buffer for 1 hr. After rinsing as above, wells were then incubated with 100 μL polyHRP streptavidin (Pierce N200) at 1:20,000 dilution in Diluent Buffer for 1 hr. After rinsing as above, wells were then incubated with 100 μL QuantaBlu Fluorogenic substrate (Pierce 15169), prepared as per manufacturer’s recommendation, for 20 min, after which 100 μL of the included Stop solution was added to each well. Plates were then read on a SpectraMax iD3 in top-read fluorescent mode at Ex 325, Em 420. Internal SpectraMax software was used to interpolate the in-well concentrations for each sample, which was exported for PK analysis. Molecules exhibited a normal biphasic distribution curve, and as such, PK parameters were determined by non-compartmental analysis for IV bolus dosing using Microsoft Excel with the PKSolver 2.0 package.

## Acknowledgments

The authors wish to thank Pauline Bariola and Jacob Felcyn for development of the SE-HPLC methodology, Jeannette Bannink for help with ELISA methodology development and instruction on PK data interpretation and analysis, Akinsola Oyelakin for help with tissue processing, Andrew J. Mhyre for *in vivo* study discussions, and Connor Burns for slide scanning. Fig. 1, A and B; Fig. 2B; Fig. 3D; Fig. 6A; Fig. 8C, and fig. S1A were created with Biorender.

## Author contributions

Z.R.C., J.P., J.M.O, and N.W.N. conceptualized the research. Z.R.C., G.P.S., P.Y., E.J.G., T-D.P., and M.H. designed the experiments. Z.R.C., G.P.S., P.Y., E.J.G., T-D.P., and M.H. performed the experiments and/or contributed to data analysis. Z.R.C. and M.H. produced figures. J.M.O. and N.W.N. contributed to funding acquisition. Z.R.C., G.P.S., E.J.G., J.M.O., and N.W.N. supervised researchers. Z.R.C., E.J.G., M.H., J.M.O., and N.W.N. wrote the manuscript. All authors contributed to review of the manuscript.

## Competing interests

Cyclera Therapeutics Inc. retains intellectual property rights to the technology described in this manuscript. Z.R.C., G.P.S., and N.W.N. own stock in and are employees of Cyclera. J.M.O. owns stock in and is an advisor of Cyclera. Z.R.C., J.M.O., and N.W.N. are inventors on patent applications for this technology. P.Y., E.J.G., T-D.P., M.H., and J.P. have no competing interests.

**Supplementary Fig. 1.**
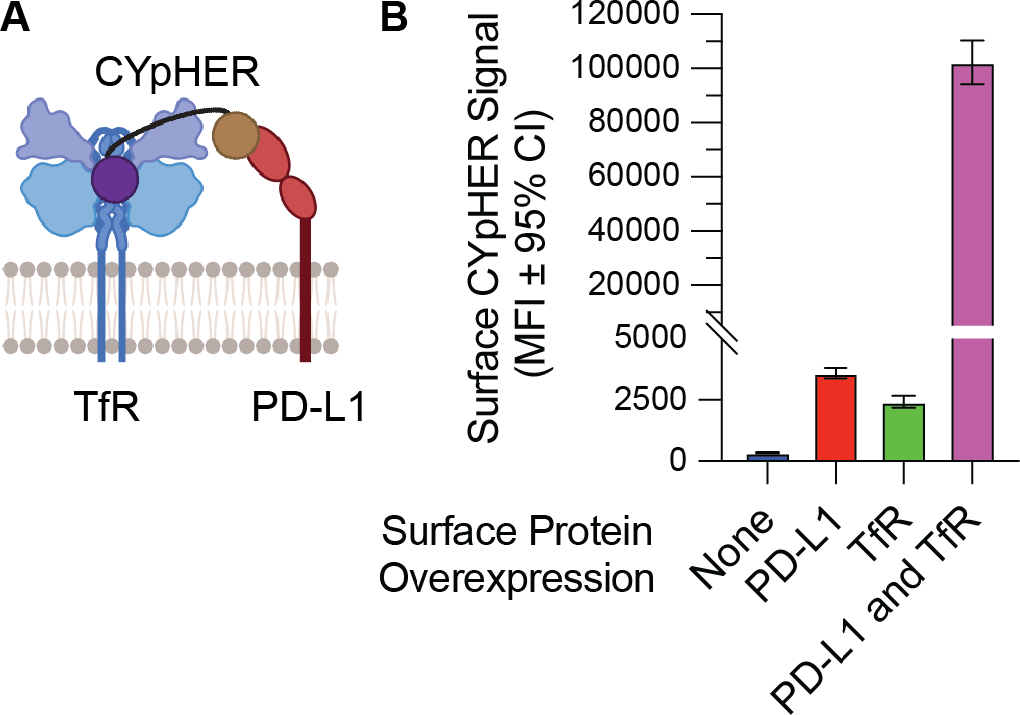
Prototype PD-L1-binding CYpHER binds both TfR and PD-L1 on cells. (**A**) Illustration of prototype CYpHER binding both TfR and PD-L1 on a cell surface. (**B**) 293T cells transfected with either TfR-RFP alone (“None” were RFP[-], “TfR” were RFP[+]) or both TfR-RFP and PD-L1-GFP (“PD-L1” were GFP[+]/RFP[-], “PD-L1 and TfR” were GFP[+]/RFP[+]) were incubated with 10 nM 6xHis-tagged prototype PD-L1 CYpHER for 24 hr and then stained with anti-His. Average anti-His signals among cells overexpressing one, the other, both, or neither of TfR and PD-L1 are shown.

**Supplementary Fig. 2.**
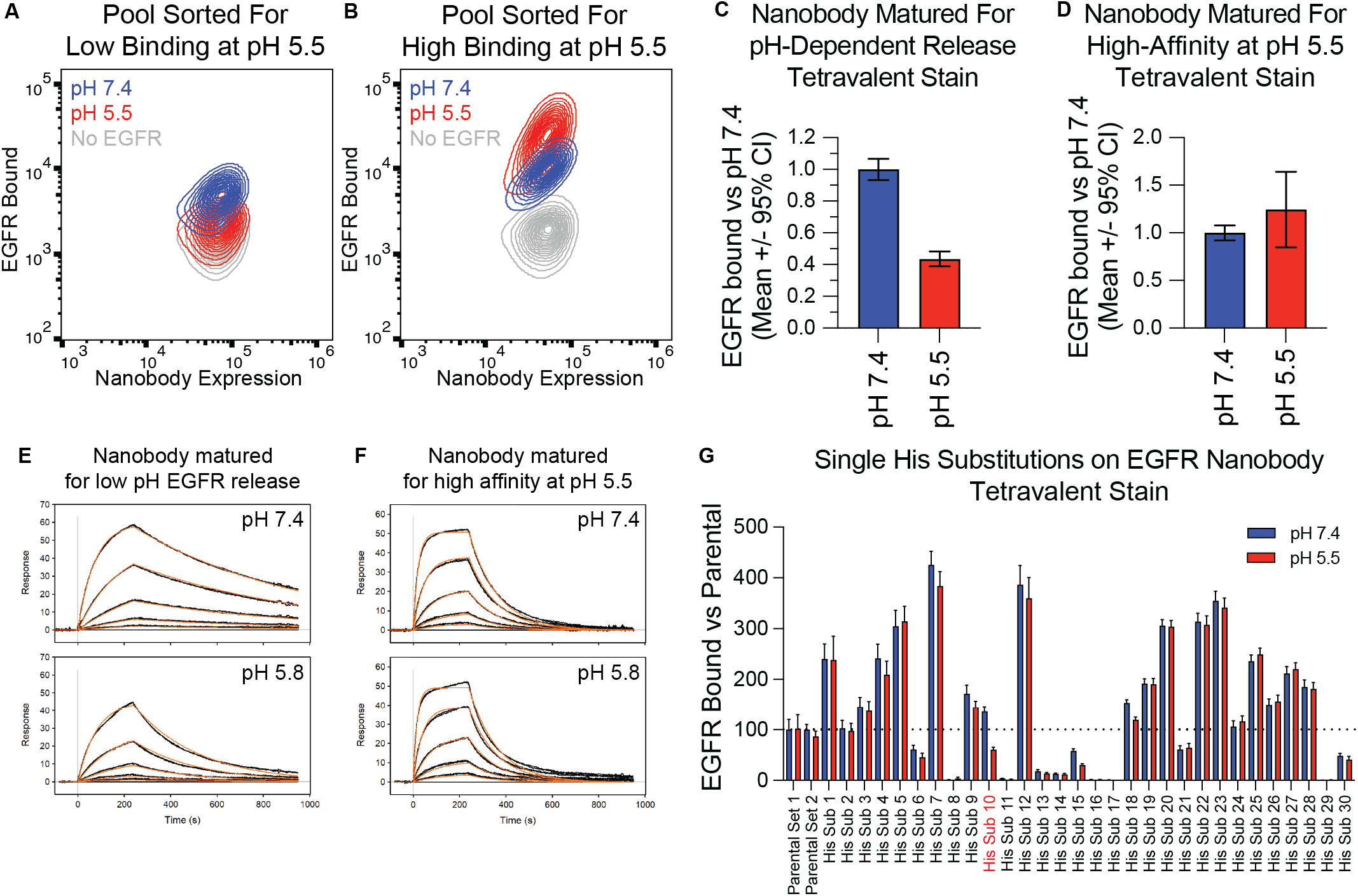
Engineering an EGFR-binding VHH nanobody for pH-dependent release. (**A** and **B**) Surface display of nanobody and stain with EGFR followed by rinse at pH 7.4 (high binding, two rounds) or pH 5.5 (low binding, two rounds) enriched for candidates with pH-dependent release. The final round of sorting at pH 5.5 was split into two populations: low binding (**A**), or high binding (**B**). (**C** and **D**) The dominant nanobody variants in the populations from (A) and (B) were stained with biotinylated EGFR and streptavidin followed by pH 7.4 or pH 5.5 rinse. The variant (**C**) from the pH 5.5 low-binding cells (**A**) lost stain in pH 5.5 vs pH 7.4, while the variant (**D**) from the pH 5.5 high-binding cells (**B**) had similar binding in both conditions. (**E** and **F**) Fc fusions of both nanobody variants were tested in surface plasmon resonance for EGFR binding in pH 7.4 (top) or pH 5.8 (bottom) buffers. (**G**) EGFR-binding VHH nanobody was subjected to conventional Histidine scanning, and singleton variants analyzed for binding to biotinylated EGFR + streptavidin followed by pH 7.4 or pH 5.5 rinse. His substitution 10 had the greatest difference in staining and was selected.

**Supplementary Fig. 3.**
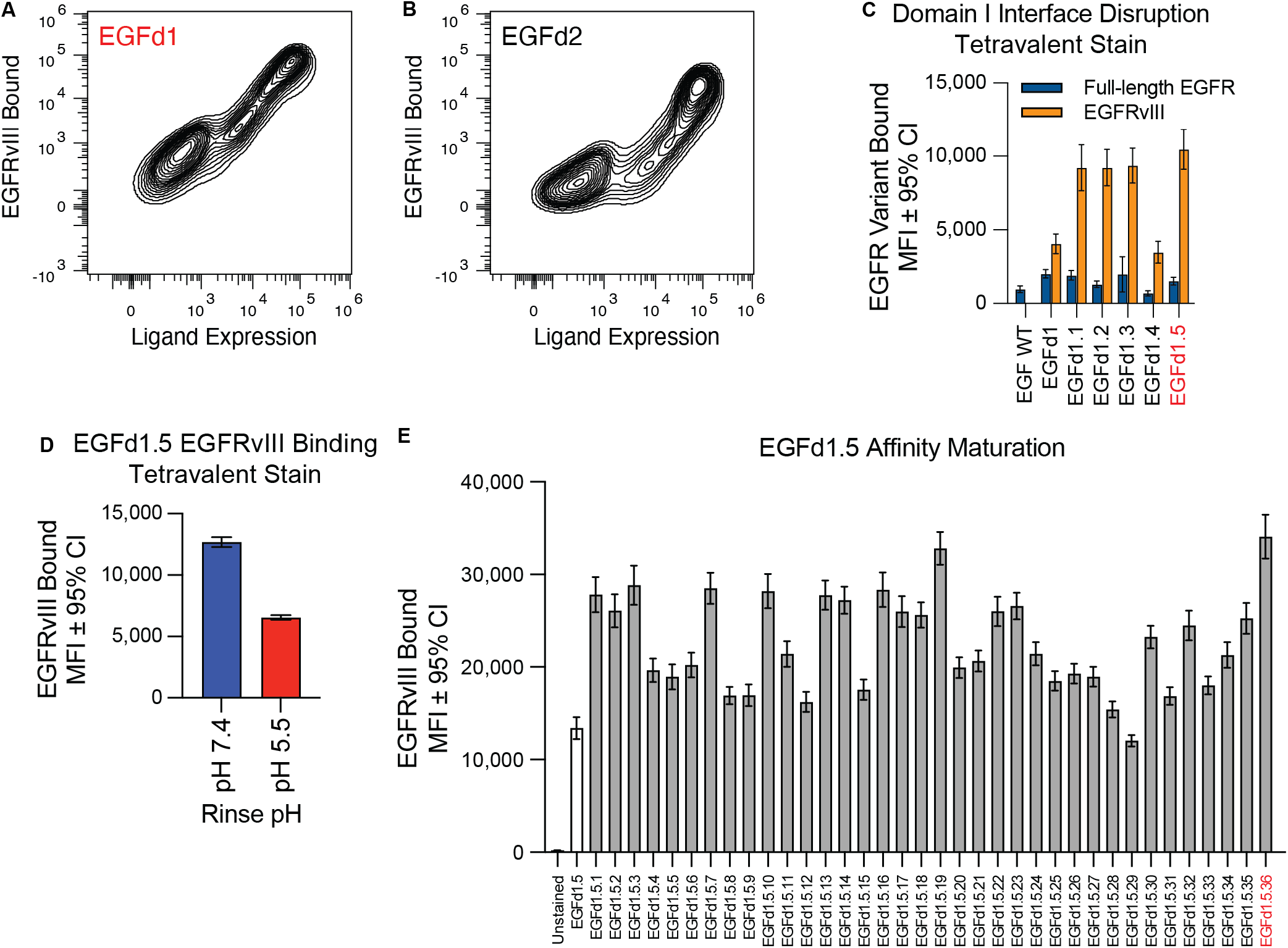
Adapting EGF for use in CYpHER. (**A** and **B**) Rosetta protein design was used with an EGF and EGFR co-crystal structure (PDB 1IVO) to design variants with improved predicted binding strength to Domain III. 1000 such variants were displayed on 293F cells and stained with biotinylated EGFRvIII and fluorescent streptavidin. After three rounds of sorting and enrichment for high-staining cells, singleton candidates were tested for EGFRvIII binding, two of which validated. One (EGFd1) was advanced. (**C**) The interface between EGF and EGFR Domain I was studied, identifying four residues predicted to be key to the interaction. These were mutated to disrupt the interaction, as singletons (EGFd1.1 to EGFd1.4) or all four at once (EGFd1.5). These were surface displayed on 293F cells and stained with biotinylated EGFR (full length) or EGFRvIII and fluorescent streptavidin. EGFd1.5 (highest staining with EGFRvIII, low binding to full length EGFR) was advanced. (**D**) EGFd1.5 was tested in surface display for biotinylated EGFRvIII + fluorescent streptavidin binding followed by pH 7.4 or pH 5.5 rinse. (**E**) Two rounds of site-saturation mutagenesis affinity maturation of EGFd1.5, followed by combining enriched mutations into 36 variant candidates, yielded binders with improved EGFRvIII binding. EGFd1.5.36 was selected.

**Supplementary Fig. 4.**
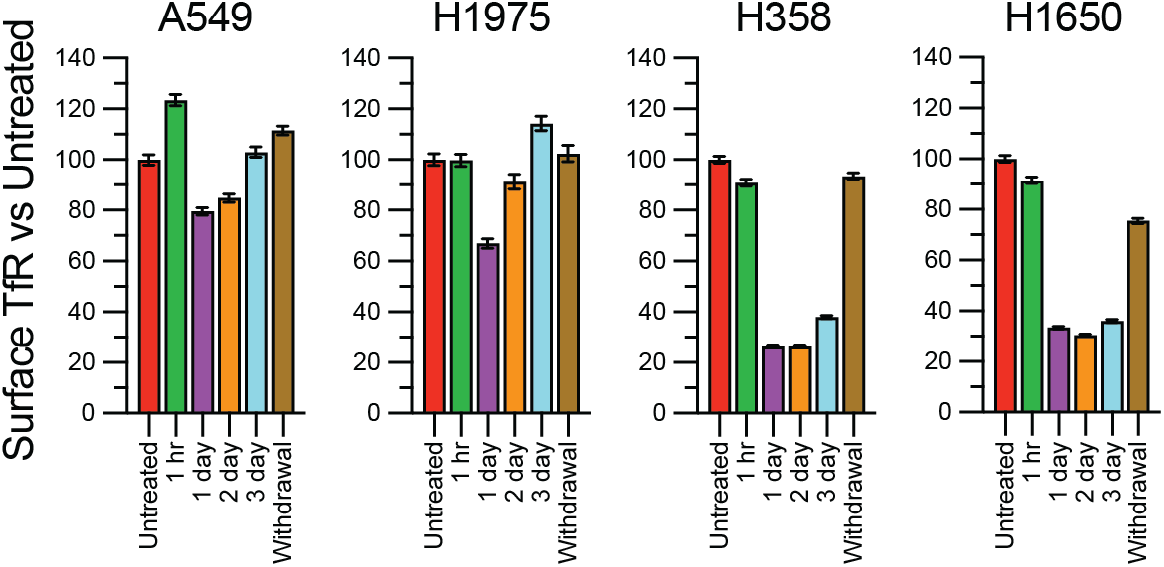
CYpHER effect on surface TfR levels in cancer cell lines. A549, H1975, H358, and H1650 cells untreated or treated with 10 nM CT-1212-1 for 1 hr, 1 day, 2 days, 3 days, or 1 day followed by 1 day without drug (“Withdrawal”) and then analyzed by flow cytometry for surface TfR levels.

**Supplementary Fig. 5.**
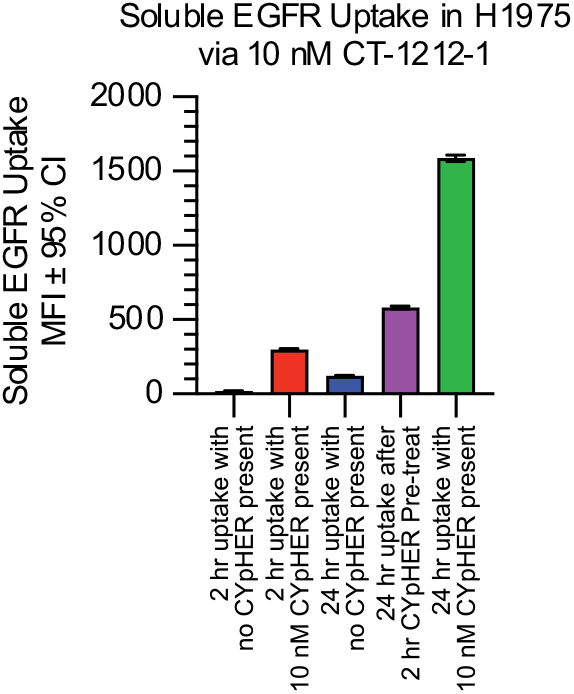
Soluble EGFR uptake with or without CYpHER withdrawal. Soluble EGFR uptake after: 2 hr with 10 nM fluorescent soluble EGFR but no CYpHER (bar 1); 2 hr with 5 nM CT-1212-1 saturated with fluorescent soluble EGFR (bar 2); 24 hr with 10 nM fluorescent soluble EGFR but no CYpHER (bar 3); 24 hr incubation with 10 nM fluorescent soluble EGFR (but no CYpHER) after 2 hr pre-treatment with 5 nM CT-1212-1 saturated with unlabeled EGFR (bar 4); or 24 hr with 5 nM CT-1212-1 saturated with fluorescent soluble EGFR (bar 5).

**Supplementary Fig. 6.**
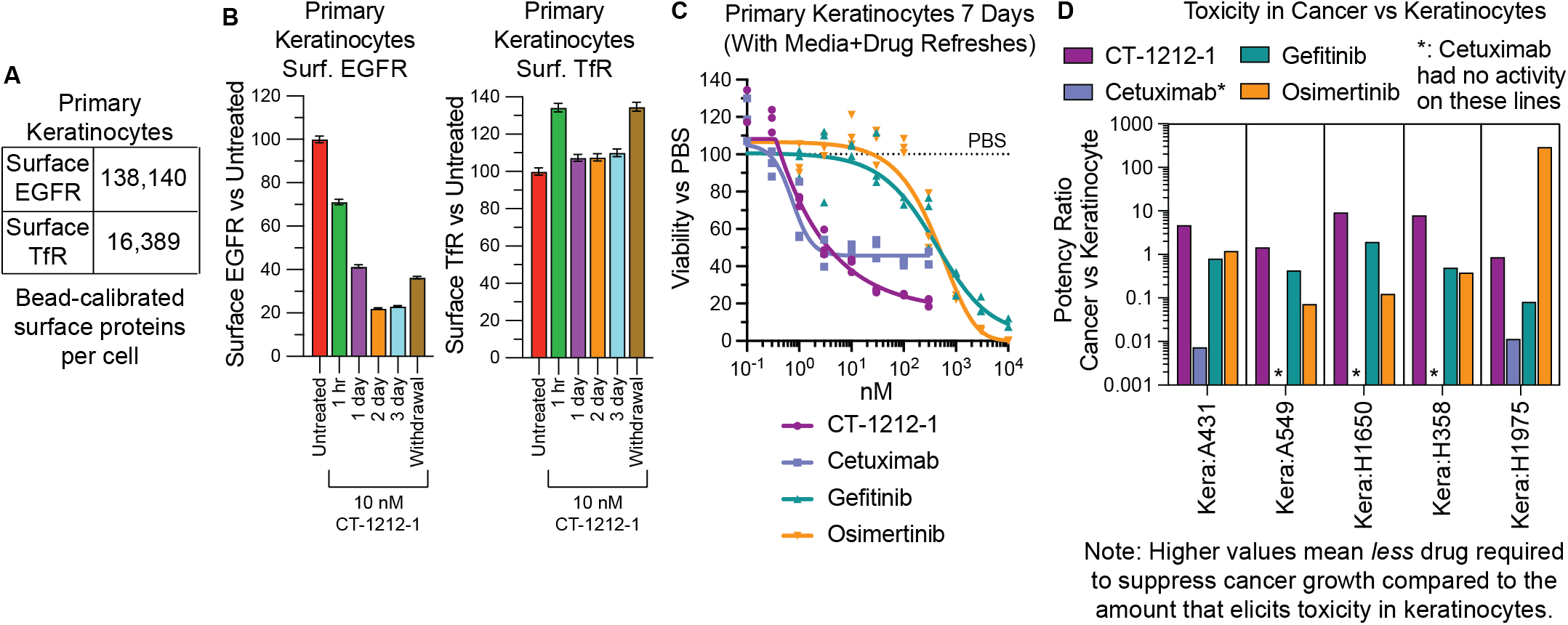
Effect of CYpHER on primary keratinocytes. (**A**) Primary human dermal keratinocytes were flow analyzed alongside calibration beads to quantitate surface EGFR and TfR levels. (**B**) Keratinocytes were untreated or treated with 10 nM CT-1212-1 for 1 hr, 1 day, 2 days, 3 days, or 1 day followed by 1 day without drug (“Withdrawal”) and then analyzed by flow cytometry for surface EGFR levels (left) and surface TfR levels (right). (**C**) Keratinocytes were treated for 7 days, with a media exchange (including drug refresh) on day 4, with CT-1212-1, cetuximab, gefitinib, or osimertinib. After treatment, cell levels per well were quantitated by CellTiter-Glo 2.0 assay. (**D**) EC50 values of cancer cell lines (Fig. 7I) compared to the EC50 values (asymmetric sigmoidal [5PL] curve fit) of the primary keratinocyte treatments. Ratio with keratinocyte EC50 as numerator and cancer line EC50 as denominator results in higher values when compound is more potent at suppressing growth of cancer lines relative to effect on keratinocyte growth.

**Supplementary Fig. 7.**
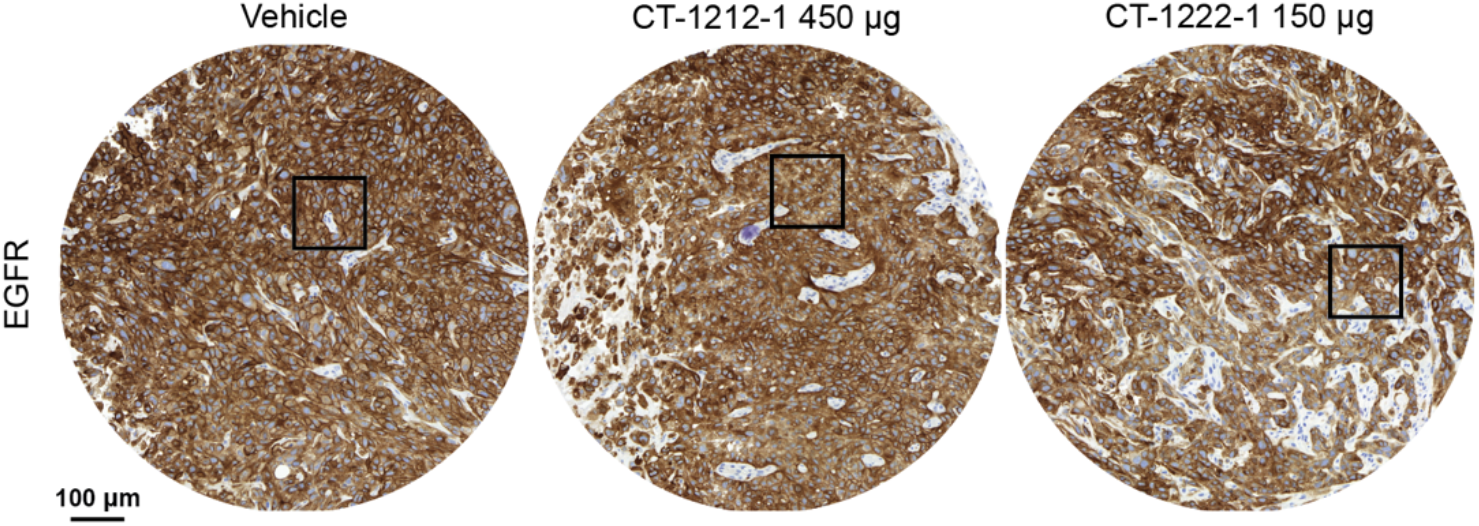
Full fields of EGFR histology shown cropped in Fig. 8F. The bounding boxes identify the crop areas used in Fig. 8F.

